# Drug metabolic activity is a critical cell-intrinsic determinant for selection of hepatocytes during long-term culture

**DOI:** 10.1101/2021.04.18.440311

**Authors:** Saeko Akiyama, Noriaki Saku, Shoko Miyata, Kenta Ite, Masashi Toyoda, Tohru Kimura, Masahiko Kuroda, Atsuko Nakazawa, Mureo Kasahara, Hidenori Nonaka, Akihide Kamiya, Tohru Kiyono, Toru Kobayshi, Yasufumi Murakami, Akihiro Umezawa

## Abstract

Hepatocytes can be used to study the pathogenesis of liver diseases and drug discovery research. Human hepatocytes are, however, hardly expandable in vitro making it difficult to secure large numbers of cells from one donor. In this study, we aimed to establish an in vitro long-term culture system that enables stable proliferation and maintenance of human hepatocytes to ensure a constant supply. We purified human hepatocytes by selection with cytocidal puromycin, and cultured them for more than 60 population doublings over a span of more than 350 days. These results show that this simple culture system with usage of the cytocidal antibiotics enables efficient hepatocyte proliferation and is an effective method for generating a stable supply of hepatocytes for drug discovery research at a significant cost reduction.

## INTRODUCTION

The liver is the largest metabolic organ in mammals and has more than 500 diverse functions, including glycogen storage, bile production, drug metabolism, ammonia metabolism, and detoxification (Trefts et al., 2017). The smallest basic unit of the liver is called a hepatic lobule, which is composed of biliary epithelial cells, hepatocytes, hepatic stellate cells, Kupffer cells, and endothelial cells. Hepatocytes are parenchymal cells that account for about 80% of the organ and are responsible for most liver functions. The morphological characteristics of hepatocytes include (1) large cell size with a diameter of about 20-30 μm, (2) round nuclei in the center of the cytoplasm, and (3) frequent multinucleation.

Hepatocytes are important as sources for cell transplantation and as target cells for gene therapy, as well as for elucidating the pathogenesis of liver diseases and for drug discovery (Dhawan et al., 2010; Ponsoda et al., 2001). Prediction of human-specific hepatotoxicity is important in drug discovery research, and inadequate predictive models can lead to drug-induced liver injury (DILI), which can cause suspension of clinical trials (Chen et al., 2015). Although laboratory animals and highly proliferative hepatocellular carcinoma cell lines are currently used, problems such as interspecies differences and variations between donors are seen in these systems. Due to the wide range of hepatic functions that must be reproduced, it is difficult to develop a suitable model system. Hepatocytes are chronically scarce due to the following problems: (1) Difficulty in securing donors of hepatocytes and (2) difficulty in *in vitro* expansion. Thus, there is a need for a highly efficient and reproducible method to prepare functional human hepatocytes.

To solve these problems, hepatocyte growth and culture systems based on liver regeneration mechanisms have been developed (Li et al., 2020; Miyajima et al., 2014). One of the characteristics of the liver is its high regenerative capacity, which is demonstrated when it is damaged by liver disease or partial hepatectomy. Among other things, hepatocytes acquire a vigorous proliferative capacity when damaged in vivo and are able to regenerate a functional liver (He et al., 2021; Li et al., 2020; Miyajima et al., 2014; Wei et al., 2021). In injured mouse livers, hepatocytes decidualized into highly proliferative hepatocytes that are characterized by (1) a phenotype that is positive for both hepatic progenitor cells and biliary epithelial markers, (2) a specific cell morphology, and (3) high proliferative capacity (Deng et al., 2018; Lu et al., 2015; Tarlow et al., 2014). Several reports suggest the presence of proliferative hepatocytes in the human diseased liver, and these cells maintain a morphologically and phenotypically intermediate state between hepatocytes and biliary epithelial cells, similar to the proliferative hepatocytes found in mice (Li et al., 2020; Schmidt and Dalhoff, 2005).

By constructing a culture system that mimics liver regeneration mechanisms, hepatocytes with proliferative potential can be induced from mouse or human mature hepatocytes in vitro (Katsuda et al., 2019, 2017; Zhang et al., 2018). Although the expression of critical markers is noticeably reduced, it has been shown that hepatocytes can be induced to mature by three-dimensional culture even after several passages. Therefore, it appears that the generation of proliferative hepatocytes using mature hepatocytes is an effective method for the efficient and stable production of a constant supply of human hepatocytes. However, the challenge still remains that these systems are unable to reproduce the vigorous proliferative capacity of human hepatocytes in vivo.

Ammonia has been used as a selection agent for enrichment of hepatocytes because of its cytotoxic effect (Tanimizu et al., 2016; Tomotsune et al., 2016; Tsuneishi et al., 2021). Puromycin has been used to select pluripotent stem cell-derived hepatocytes with a high expression of CYP3A4 (Miyata et al., 2020). Puromycin is an aminonucleoside antibiotic produced by Streptomyces alboniger (Aviner, 2020; Miyamoto-Sato et al., 2000; Yarmolinsky and Haba, 1959). Puromycin is commonly used as a selection compound for genetically modified cell lines and is useful as a probe for protein synthesis. Its structure is similar to that of the 3’ end of aminoacyl-tRNA, and it can enter the A site of the ribosome and bind to the elongating strand, thereby inhibiting protein synthesis. This reaction, called puromycinylation, is energy-independent and causes degradation of the 80S ribosome (Aviner, 2020). The inhibition of protein synthesis is non-specific and is a result of competition with aminoacyl-tRNA (Miyamoto-Sato et al., 2000).

There is a shortage of human hepatocytes for drug discovery research and cell transplantation, and their widespread use requires a culture system capable of efficient, large-scale production of hepatocytes that maintain their functionality. The purpose of this study was to establish a long-term in vitro culture system that enables stable proliferation and maintenance of functional human hepatocytes with the following three features: (1) Establishment of cells with the characteristics of proliferative hepatocytes, (2) maintenance of proliferation and functionality for a long period of time in vitro, and (3) induction of hepatic maturation by three-dimensional culture. We established a culture system that enables simple and efficient hepatocyte proliferation and is an effective method for a stable supply of hepatocytes for drug discovery research and cell transplantation.

## RESULTS

### Propagation of human hepatocytes

We first investigated whether human hepatocytes were expandable in vitro. We isolated hepatocytes from a DILI patient’s liver (#2064) and assessed the proliferative capacity (Figure 1A). Hepatocytes formed colonies a few days after seeding (Figure S1A) and proliferated until they reached confluence. After each passage, proliferating human hepatocytes (ProliHH) exhibited colony-like morphology and continued growing until they became confluent (Figure 1B). ProliHH maintained their proliferative capacity for up to 200 days and proliferated nearly 10^13^-fold for 21 passages (Figure 1C and 1D). ProliHH were small in size and exhibited a high nucleus-to-cytoplasm ratio in early passages (~P6) (Figure S1B), but the nuclear-cytoplasmic ratio of proliHH became similar to that of PHH after several passages. The volume of hepatocytic cytoplasm increased over time and the cells stopped proliferating after 21 passages (Figure 1B). Immunocytochemical analysis revealed that ProliHH at P5 was positive for a proliferation marker Ki67, but most of cells were negative at P21 (Figure 1E). On the other hand, there were SA-β-gal positive cells at P21 but not P5 (Figure 1F). Karyotypic analysis revealed that ProliHH at P13 with high proliferative capacity maintained a normal diploid karyotype, 46XX (Figure 1G).PHH stably proliferated at least to P21 without chromosomal abnormality by co-culturing with mouse fetal fibroblasts

**Figure 1.**
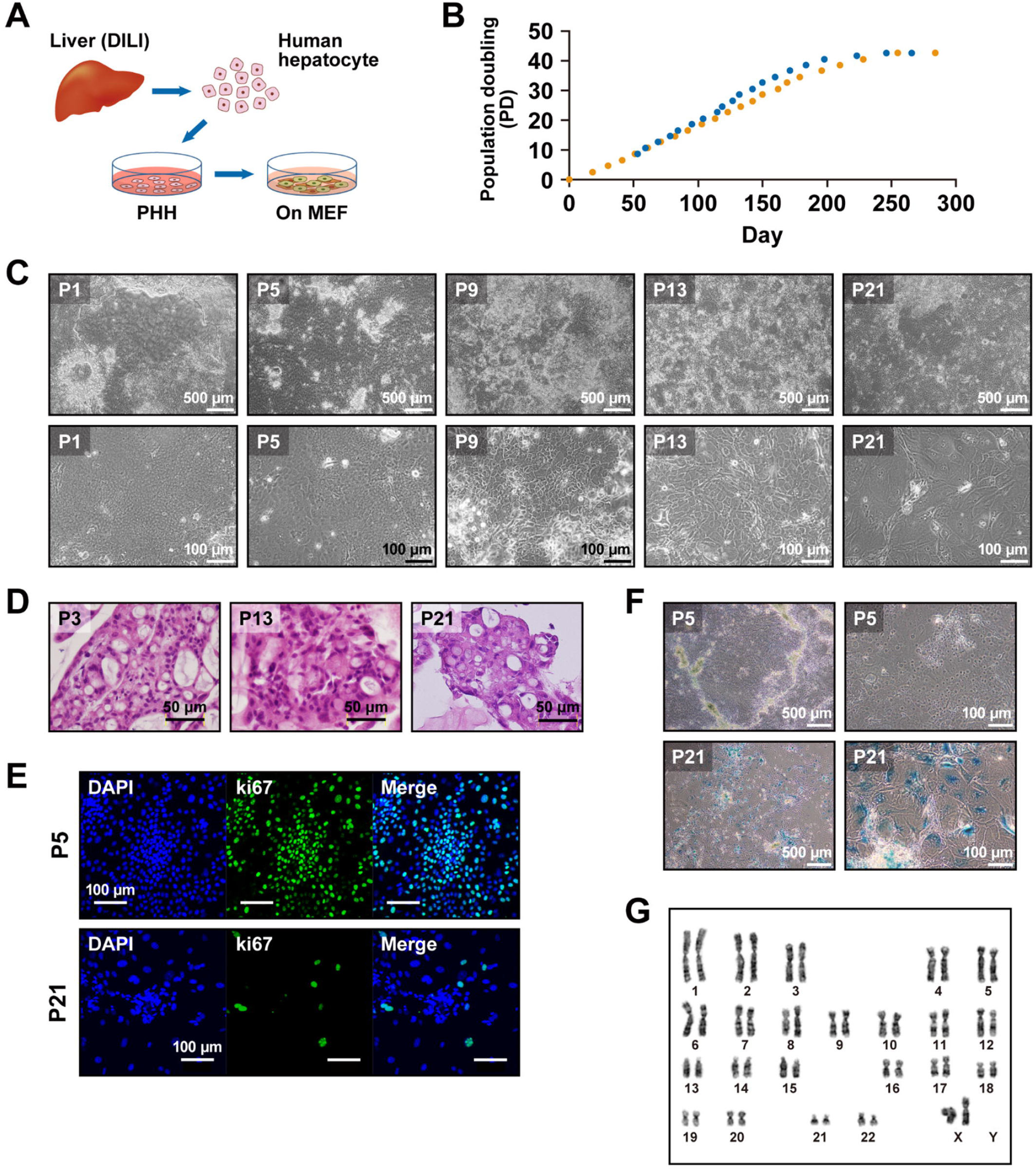
Establishment of a long-term hepatocyte culture. A. Scheme of culture protocol. B. Phase-contrast photomicrographs of ProliHH from passage 1, 5, 9, 13, and 21. C. Growth curves of ProliHH. Proliferative capacity was analyzed at each passage. Cells were passaged in the ratio of 1:4 for each passage (n=2). “Population doubling” indicates the cumulative number of divisions of the cell population. D. Microscopic view of ProliHH at passages 3, 13, and 21. HE stain. E. Immunocytochemical analysis of ProliHH with an antibody to Ki67 (cell proliferation marker). F. A senescence-associated beta-galactosidase stain of ProliHH at the indicated passages. The number of β-galactosidase-positive senescent cells increased at passage 21. G. Karyotypes of ProliHH at passage 13. Hepatocytes had 46XX and did not exhibit any significant abnormalities.

To characterize the in vitro proliferating human hepatocytes, ProliHH, the expression of hepatocyte and biliary epithelial cell (BEC) markers was analyzed. ProliHH expressed hepatocyte-associated genes such as ALB and AAT (Figure 2A) and showed decreased expression of the genes for ALB and cytochrome P450 (CYP) 1A2. In contrast, ProliHH showed increased expression of AAT and CYP3A4 during the first few passages and decreased expression at later stages. The expression of CK7 and CK19 varied depending on the number of passages (Figure 2A, S1C, and S1D). Immunocytochemistry revealed that ProliHH expressed the hepatocyte marker ALB and the BEC marker CK7 at both early and late passages (P5 and P21) (Figure 2B and 2D). The hepatocyte marker HNF4A was positive in an early passage (P5) but was later negative (P21) (Figure 2C). Also, binuclear cells were detected at P21 (Figure 2D, circles). These results indicate that ProliHH had both biliary and hepatic characteristics and maintained a high proliferative capability for more than 200 days. Glycogen storage capacity was also observed at P21, when ProliHH stopped dividing, but not at P3 and P13 (Figure 2E and 2F). We then evaluated ProliHH for CYP induction. We investigated expression levels of the three major CYP enzymes, CYP1A2, CYP2B6, and CYP3A4, in early and middle passages (P5 and P11) (Figure 2G, H, I). We exposed the cells to omeprazole for 24 h, phenobarbital for 48 h, and rifampicin for 48 h. Expression of CYP1A2 was up-regulated 21-fold upon exposure to omeprazole (Figure 2G); CYP2B6 expression was not induced upon exposure to phenobarbital (Figure 2H); and expression of CYP3A4 was up-regulated 1.9- and 2.4-fold after exposure to rifampicin at P5 and P11, respectively (Figure 2I). CYP3A4 activity was higher than HepG2 in an early passage (P5), but decreased with passage number (Figure 2J).

**Figure 2.**
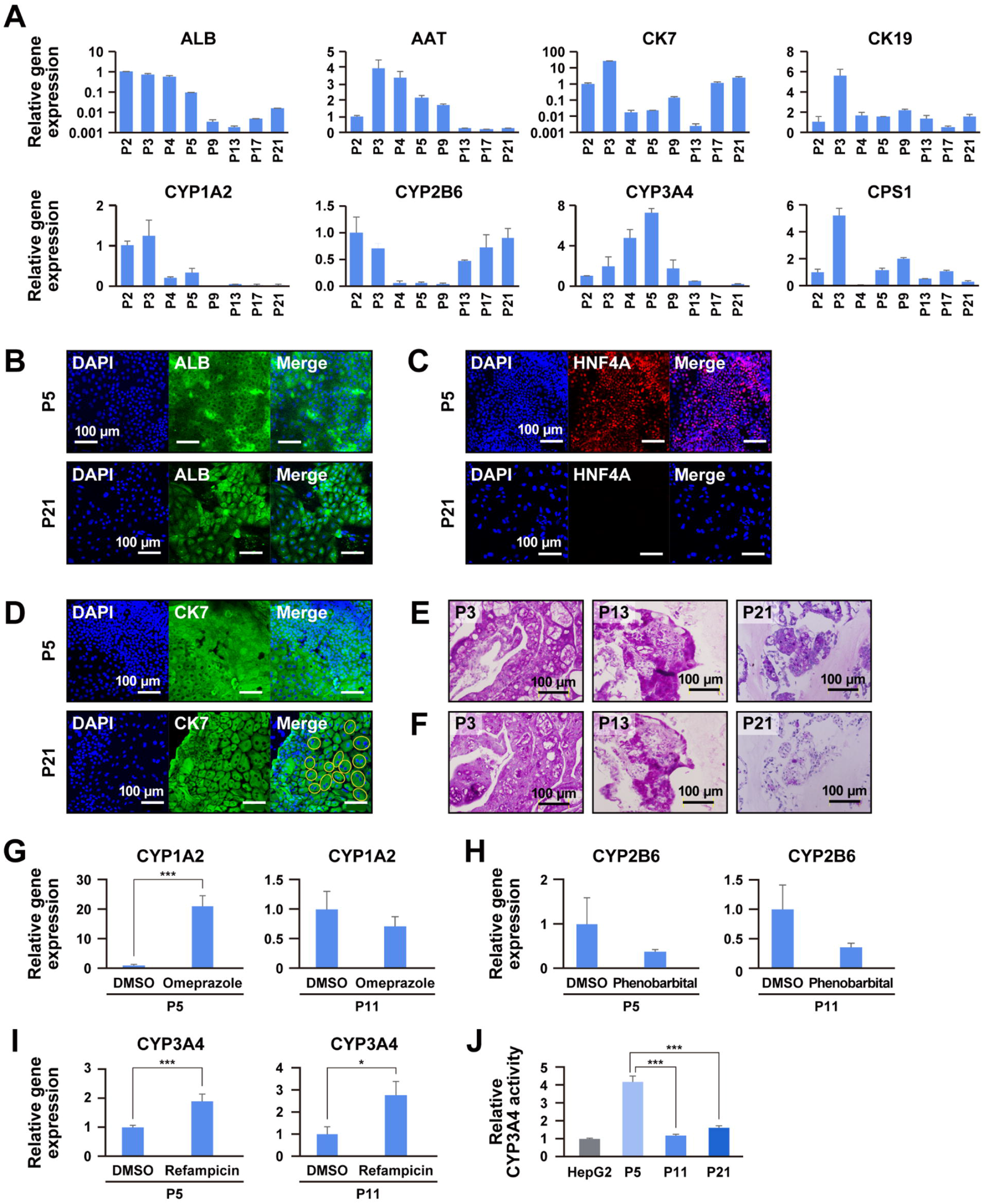
Characteristic analysis of ProliHH during long-term culture. A. Gene expression of ProliHH from passage 2 to 21 by qRT-PCR. Data were normalized with the housekeeping UBC gene. The expression level of each gene in passage 2 was set to 1.0. Error bars indicate the standard deviation (n=3). B-D. Immunocytochemical analysis of ProliHH with antibodies to ALB (B), HNF4A (C), and bile duct marker CK7 (D) at the early and late passages (passage 5 and 21, respectively). Yellow circles indicate binuclear cells (D). E-F. Glycogen storage in hepatocytes (passage 3, 13, and 21) by PAS stain with (E) and without (F) diastase digestion. G-I. Expression of the genes for CYP1A2 (G), CYP2B6 (H), and CYP3A4 (I) in hepatocytes after exposure to omeprazole (G), phenobarbital (H), or rifampicin (I). The expression level of each gene without any treatment (DMSO) was set to 1.0. Each expression level was calculated from the results of independent (biological) triplicate experiments. Error bars indicate standard error. The student’s T-test was performed for statistical analysis of two groups. *p < 0.05, **p < 0.01, ***p < 0.001. J. CYP3A4 activity of hepatocytes (passage 5, 11, and 21). The CYP3A4 activity of HepG2 cells was set to 1.0. Error bars represent the standard errors. Each expression level was calculated from the results of independent (biological) triplicate experiments. Error bars indicate the standard error. ***p < 0.001.

### Selection of proliferative hepatocytes by puromycin treatment

Hepatocytes have drug-metabolizing activity, including CYP3A4, which is responsible for detoxification and metabolism of antibiotics. We therefore exposed ProliHH to different concentrations of puromycin for 3 days to determine if the ProliHH were resistant (Figure 3A). The ProliHH showed resistance to puromycin at concentrations ranging from 1 μg/mL to 100 μg/mL (Figure 3B), while MEF did not (Figure S2A). ProliHH selected with puromycin were small and displayed a high nuclear-to-cytoplasmic ratio with clear nucleoli (Figure 3C and S2B). We then performed puromycin treatment on ProliHH from other DILI patients (donor ID: 2061 and 2062) to determine if these results were reproducible (Figure S2C, S2D, and S2E). The puromycin-selected ProliHH (donor ID: 2061 and 2062) exhibited essentially the same morphology as ProliHH (donor ID: 2064). Gene expression analysis revealed that exposure to puromycin suppressed the expression of mesenchymal cell markers (COL1A1 and ASMA) but enhanced hepatocyte markers (ALB and AAT), cytochrome P450 genes (CYP1A2, CYP2B6, CYP3A4, and CYP2C9), and hepatic progenitor-associated markers (CK7, CK19, EpCAM, SOX9, and PROM1) (Figure S2F).

**Figure 3.**
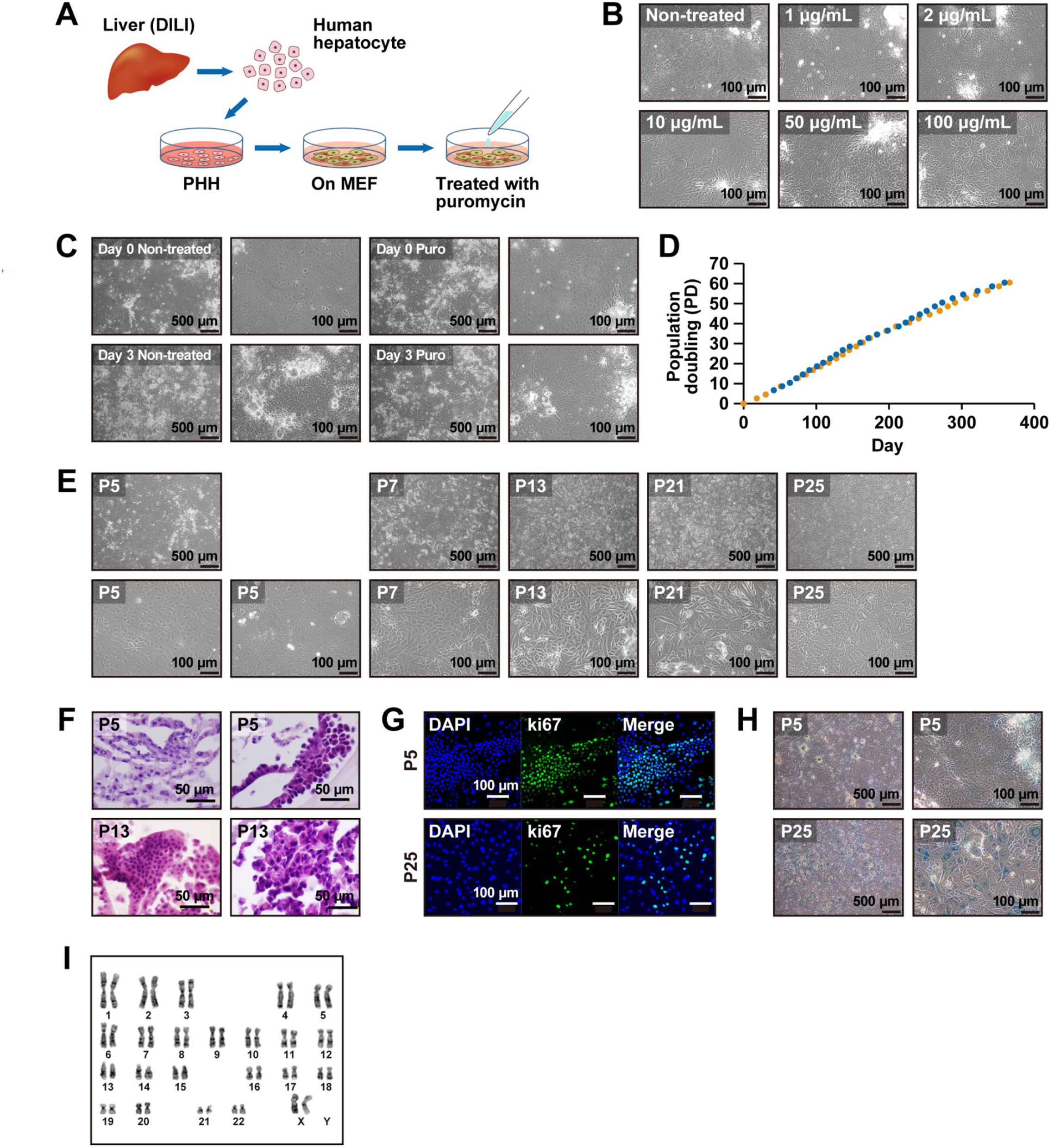
Puromycin-based selection of ProliHH. A. Scheme of culture protocol. Puromycin was added at each passage for selection. B. Phase-contrast photomicrographs of ProliHH with exposure of puromycin for 3 days. Different concentrations of puromycin (0, 1, 2, 10, 50 and 100 μg/mL) were added at 80% confluence. C. Phase-contrast photomicrographs of ProliHH cells before and after 2 μg/mL puromycin. Puromycin was added when the cells reached confluence (Day 0) and removed 3 days after the addition (Day 3). D. Growth curves of puromycin-treated ProliHH in duplicated experiments. Proliferative capacity was analyzed at each passage. Cells were passaged in the ratio of 1:4 at each passage (n=2). “Population doubling” indicates the cumulative number of divisions of the cell population. E. Phase-contrast photomicrographs of puromycin-treated ProliHH at passage 5, 7, 13, 21, and 25. F. Microscopic view of puromycin-treated ProliHH at passage 5, 13, and 25. HE stain. G. Immunocytochemical analysis of puromycin-treated ProliHH with an antibody to Ki67 (cell proliferation marker). H. A senescence-associated beta-galactosidase stain of puromycin-treated ProliHH at passages 5 and 25. The number of β-galactosidase-positive senescent cells increased at passage 25. I. Karyotypes of puromycin-treated ProliHH at Passage 24. Details are given in Supplemental Figure S3B.

Next, we assessed the proliferative capacity of puromycin-treated cells (Figure 3D). The cells continued to proliferate to 60 population doublings over more than 350 days, resulting in a 10^18^-fold increase in cell number. The proliferating cells were small, with a high nucleus-to-cytoplasm ratio in the early passages (~P6); but after P20, the ratio of nuclei to cytoplasm was similar to that of hepatocytes. The cells increased in homogeneity during propagation from P5 to P20, and no senescence-like morphology such as significant cytoplasmic enlargement was observed (Figure 3E and 3F). Early on (P5), the cells were positive for the proliferative marker Ki67, but the number of Ki67-positive cells decreased in later passages (P25) (Figure 3G). By P25 the cells were positive for SA-β-gal, a senescence marker (Figure 3H). Karyotypic analysis showed that at P24, the cells had maintained a normal karyotype, 46XX (Figure 3I).

We examined the puromycin-selected ProliHH after long-term culture for expression of hepatocyte- and BEC markers. The cells continued to express genes for hepatocyte and bile duct markers (Figure 4A and S3A). The expression of cytochrome P450 enzymes, CYP1A2, CYP2B6, and CYP3A4, in puromycin-treated cells at P4 was higher compared to non-treated cells (Figure 4B). Hepatocyte markers such as ALB and AAT were also upregulated in puromycin-treated cells at P11 and P17. In contrast, the expression levels of mesenchymal cell markers were significantly decreased in puromycin-treated cells, probably due to the elimination of mesenchymal cells and the selection of puromycin-resistant cells (Figure 4B).

**Figure 4.**
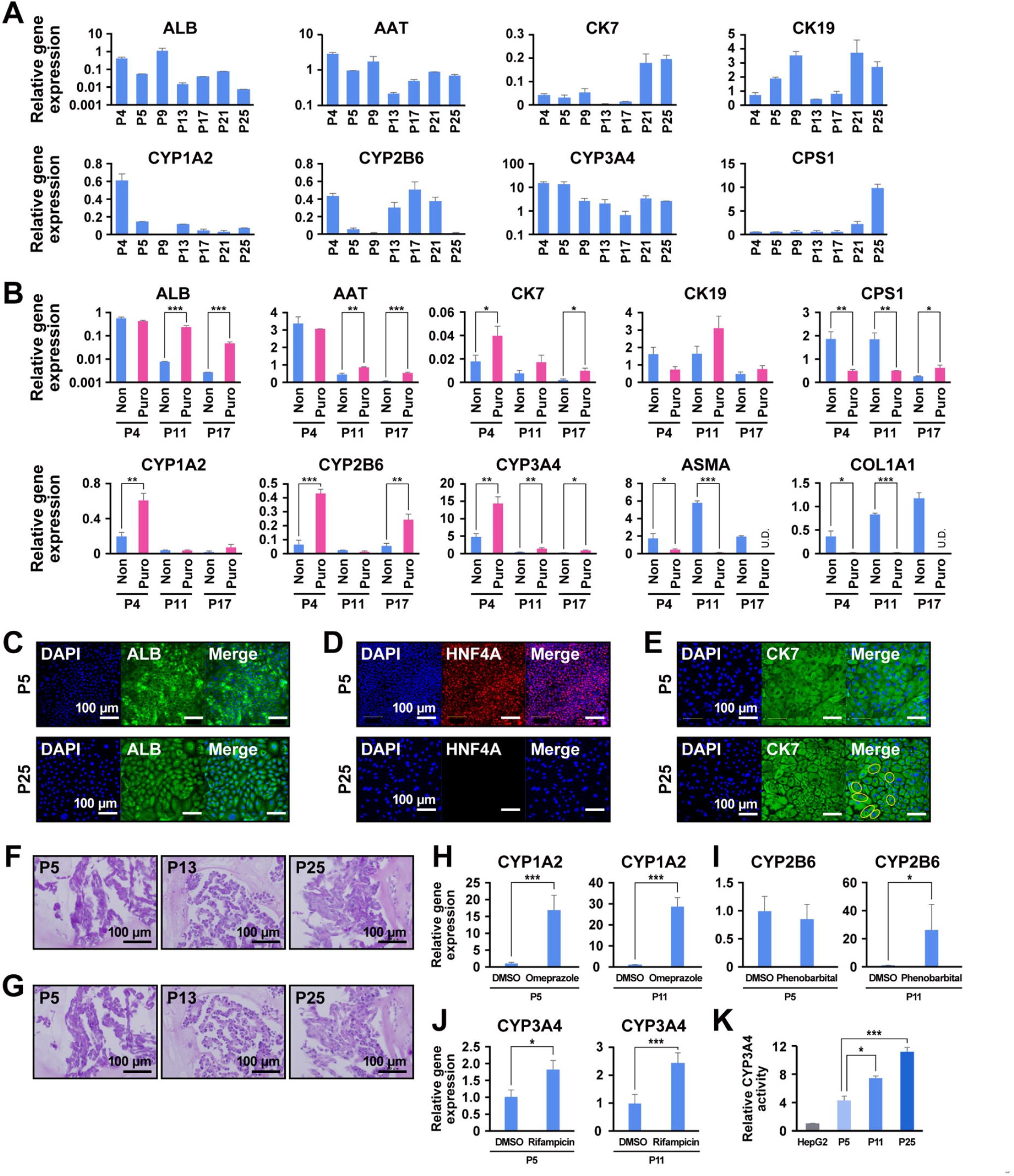
Characteristic analysis of puromycin-selected hepatocytes during long-term culture. A. Gene expression by qRT-PCR in puromycin-treated ProliHH from passage 4 to 25. Data were normalized with the housekeeping gene UBC. The expression level of each gene at passage 2 was set to 1.0. Error bars indicate the standard deviation (n=3). B. Gene expression by qRT-PCR in puromycin-treated ProliHH at passages 4, 11, and 17 by qRT-PCR. Puromycin-treated (“Puro”) and non-treated (“Non”) ProliHH at passages 4, 11, and 17 were compared. The data were normalized by the housekeeping UBC gene. The expression level of each gene in passage 2 was set to 1.0. Error bars indicate the standard deviation. Each expression level was calculated from the results of independent (biological) triplicate experiments. The student’s T-test was performed for statistical analysis of two groups. *p < 0.05, **p < 0.01, ***p < 0.001. C-E. Immunocytochemical analysis of puromycin-treated ProliHH with antibodies to ALB (C), HNF4A (D), and bile duct marker CK7 (E) at the early and late passages (passage 5 and 25, respectively). Yellow circles indicate binuclear cells (E). F-G. Glycogen storage in puromycin-treated hepatocytes (passage 5, 13, and 25). (F) PAS stain. (G) PAS stain with diastase digestion. H-J. Expression of the genes for CYP1A2 (H), CYP2B6 (I), and CYP3A4 (J) in puromycin-treated hepatocytes after exposure to omeprazole (H), phenobarbital (I), or rifampicin (J). The expression level of each gene without any treatment (DMSO) was set to 1.0. Each expression level was calculated from the results of independent (biological) triplicate experiments. Error bars indicate standard error (n=3). The student’s T-test was performed for statistical analysis of two groups. *p < 0.05, ***p < 0.001. K. CYP3A4 activity of hepatocytes (passage 5, 11, and 25). The CYP3A4 activity of HepG2 cells was set to 1.0. Error bars represent standard errors (n=3). Each expression level was calculated from the results of independent (biological) triplicate experiments. Error bars indicate the standard error. *p < 0.05, ***p < 0.001.

Immunocytochemistry revealed that the cells expressed ALB and CK7 in both early and late passages (P5 and P25, respectively) (Figure 4C and 4E). The hepatocyte marker HNF4A was positive at an early passage (P5), whereas it was negative at a later passage (P25) (Figure 4D). With puromycin treatment, the number of binuclear ProliHH were increased at P25 (Figure 4E). Glycogen storage was not detected by PAS staining in ProliHH that had been treated with puromycin (Figure 4F and 4G). CYP1A2 and CYP3A4 were significantly upregulated by omeprazole and rifampicin (Figure 4H, I, J). Furthermore, the cells continuously exposed to puromycin increased CYP3A4 activity (Figure 4K). We performed the same experiments with lower concentrations (1 μg/mL) of puromycin and obtained the same results regarding morphology, proliferation, and gene expression, and maintained CYP3A4 activity after passaging (Figure S3C-H and S4).

### Global gene expression analysis

Gene expression profiles of in vitro proliferating hepatocytes; ProliHH, were compared to primary human hepatocytes (PHH) from DILI patients, non-cultured fresh mature hepatocytes (fresh MH), iPSC-derived hepatocyte-like cells and bile duct epithelial cells (Figure 5). Principal component analysis and hierarchical clustering analysis revealed that “non-treated” and “puromycin-treated” cells were clustered into independent groups, regardless of the passage number (Figure 5A). A heatmap showed that the cells have both hepatocyte and BEC characteristics at all passages (Figure 5B, 5C). To further elucidate differences between the groups, we identified differentially expressed genes in PHH, puromycin-treated and non-treated ProliHH (Figure 5D). Compared with PHH, 3186 genes were significantly up-regulated in ProliHH and 3551 genes in puromycin-treated ProliHH, of which 3068 (83.6%) were coincidentally up-regulated (Figure 5E). Over-Representation Analysis was performed on the 3068 genes to identify the pathways that were significantly related to the gene expression (Figure 5E). We found that the ProliHH retained a hepatocyte-related gene expression pattern. Puromycin-treated and non-treated ProliHH showed enhanced expression of genes related to hepatocyte functions such as fatty acid metabolism, drug metabolism, amino acid degradation, and ammonium metabolism. We also identified enhanced expression of genes related to ERBB signaling pathways, which play an important role in liver regeneration and hepatocyte proliferation.

**Figure 5.**
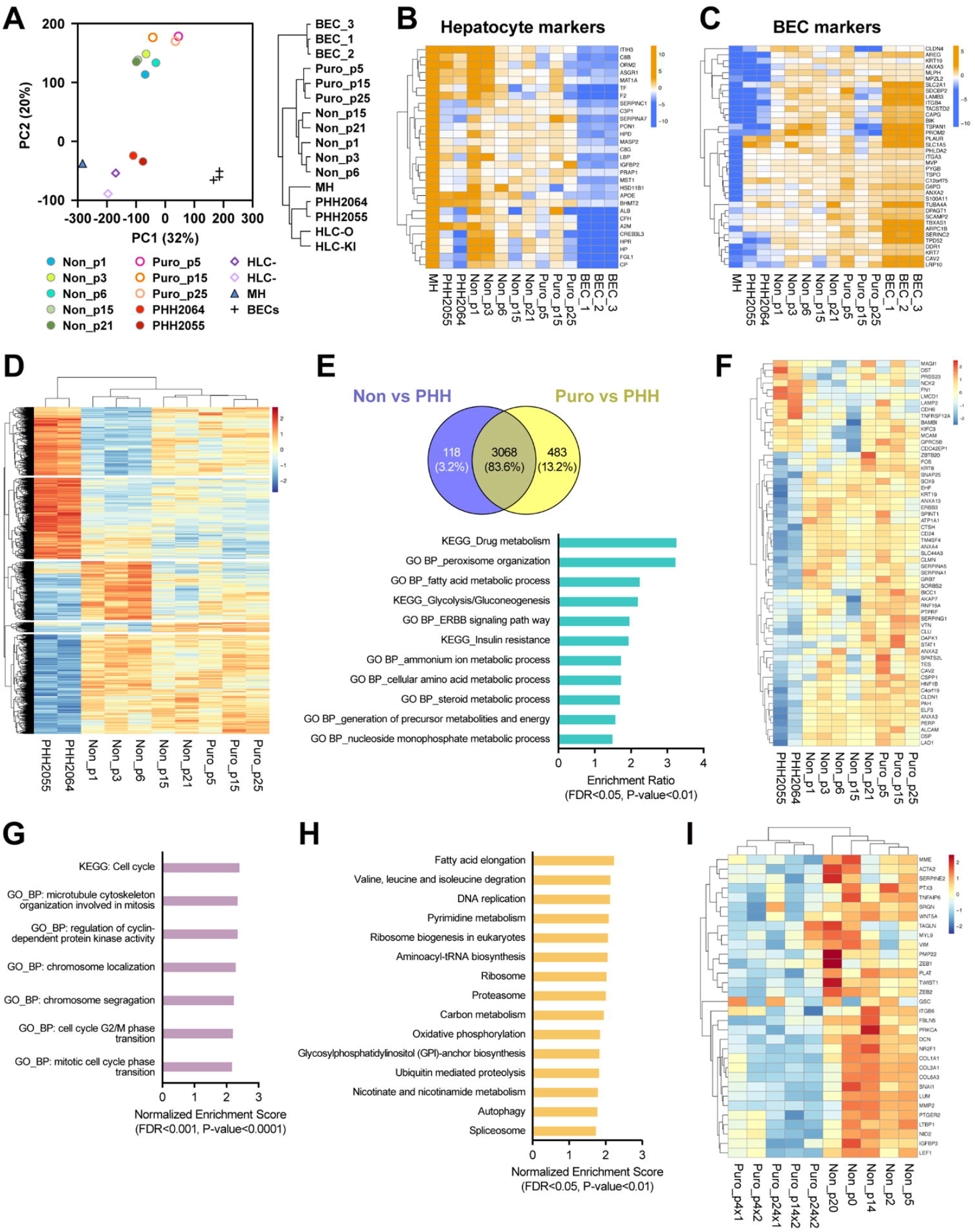
Global gene expression analysis of hepatocytes reveals a distinct hepatocyte group with proliferative capability. A. Principal-component analysis (PCA) for non-treated ProliHH (Non) at passage 1, 3, 6, 15, and 21), puromycin-treated ProliHH (Puro) at passage 5, 15, and 25, PHH (PHH2064 and PHH2055), mature hepatocytes (MH), biliary epithelial cells (BEC: GSM4454532, GSM4454533, GSM4454534), and iPSC-derived hepatocyte-like cells (HLC-O and HLC-KI) by global gene expression. Right: Hierarchical clustering analysis of expression profiles of the samples shown in PCA. B-C. Heatmaps of the liver-associated genes. Genes upregulated in mature hepatocytes (B) and bile duct epithelial cells (C) were used for the heatmap analysis (gene list: https://www.nature.com/articles/s41467-019-11266-x). The average gene expression in non-treated and puromycin-treated ProliHH was set to 0. Their expression levels were compared in groups of mature hepatocytes (MH), primary hepatocytes (PHH), and bile duct epithelial cells (BEC). The color bar indicates the signal intensity at log2 expression. D. A heatmap for differentially expressed genes (1.5 < fold changes and p < 0.05) in the comparisons of PHH versus non-treated ProliHH versus puromycin-treated ProliHH. The color bars show the signal strength scaled by the z score. E. Up-regulated genes (1.5 < fold changes and p < 0.05) in non-treated and puromycin-treated ProliHH compared to PHH. Among the genes identified, 3,068 genes (83.6%) matched. Bar graph: Over-Representation Analysis (ORA) of 3,068 genes with WebGestalt (WEB-based Gene SeT AnaLysis Toolkit) (FDR < 0.05, P < 0.01). F. Heatmap showing the expression levels of top fetal hepatobiliary hybrid progenitor up-regulated genes (gene list: https://www.nature.com/articles/s41467-019-11266-x) in PHH, non-treated and puromycin-treated ProliHH. The color bars show the signal strength scaled by the z score. G. Gene Set Enrichment Analysis was performed to identify enriched gene sets in puromycin-treated ProliHH compared to non-treated ProliHH (FDR < 0.0001, p-value < 0.0001). H. The top 15 genesets enriched in puromycin-treated hepatocytes. Gene sets that were significantly upregulated in puromycin-treated ProliHH compared to non-treated ProliHH were clustered by affinity propagation method (FDR < 0.05, P < 0.01) and ranked based on the normalized enrichment score1 (NES). I. Heatmap of genes with upregulated expression during epithelial-mesenchymal transition in non-treated and puromycin-treated ProliHH (gene list: https://journals.plos.org/plosone/article?id=10.1371/journal.pone.0051136). The colored bars show the signal strength in scaled by z score. “Non”: non-treated ProliHH, “Puro_x1’’: ProliHH treated with 1 μg/ml puromycin, “Puro_x2’’: ProliHH treated with 2 μg/ml puromycin.

Puromycin-treated and non-treated ProliHH exhibited significantly enhanced expression of fetal hepatobiliary hybrid progenitor and hepatic progenitor-related genes such as AFP, SOX9, PROM1, and EpCAM (Figure 5F, S5A, S5B, and S5C). Together, these results suggest that puromycin-treated and non-treated ProliHH share common characteristics with hepatic progenitors. To elucidate the effect of puromycin, we compared gene expression of puromycin-treated and non-treated cells and identified pathways by Gene Set Enrichment Analysis. The results show that the enrichment of gene sets related to cell proliferation and division was much higher (FDR < 0.0001, p-value < 0.0001) in puromycin-treated ProliHH than in non-treated cells (Figure 5G). Puromycin-treated ProliHH maintain a high proliferative capacity in a longer time-period, compared with the non-treated cells. The results of Gene Set Enrichment Analysis and the cell growth are exactly consistent. In addition to the DNA replication pathway, the amino acid degradation pathway, the fatty chain elongation pathway, and the oxidative phosphorylation pathway were identified as “enhanced pathways” (Figure 5H and S5D). Notably, the biosynthesis of ribosomes and tRNA, which is the active site of puromycin, was enhanced. This may indicate a homeostatic response of the cells to puromycin treatment. The epithelial-mesenchymal transition core genes were enriched in non-treated ProliHH (Figure 5I and S5E).

### Hepatocyte maturation

We next investigated whether puromycin-treated and non-treated ProliHH could acquire reversible mature hepatocyte properties. Puromycin-treated and non-treated ProliHH at early (P5), middle (P11), and late (P21 or P25) passages were cultured in low-attachment plates for 10 days for three-dimensional (3D) culture (Figure 6A and S6A). Under 3D-culture conditions, puromycin-treated and non-treated ProliHH formed spheroids irrespective of the number of passages (Figure 6B-6D). Glycogen storage, which could not be detected in two-dimensional culture of puromycin-treated and non-treated ProliHH, was confirmed in 3D culture (Figure 6E-6J). Hepatocyte markers (ALB, CYP3A4, and MRP2) were expressed in 3D-cultured puromycin-treated and non-treated ProliHH (Figure 6K-6M). Expression levels of hepatocyte markers, especially ALB, AAT, CYP1A2, CYP2B6, and CYP3A4, were significantly increased with maturation in 3D culture, whereas the expression of CK7, BEC and hepatocyte progenitor marker, was substantially suppressed (Figure 7A, 7B and 7C). It is noteworthy that maturation of puromycin-treated ProliHH was observed in both early and late passages of ProliHH (Figure 7, S6 and S7). These results indicate that puromycin-treated ProliHH is highly capable of regaining characteristics similar to mature hepatocytes, suggesting that the addition of puromycin is effective in maintaining the characteristics of mature hepatocytes.

**Figure 6.**
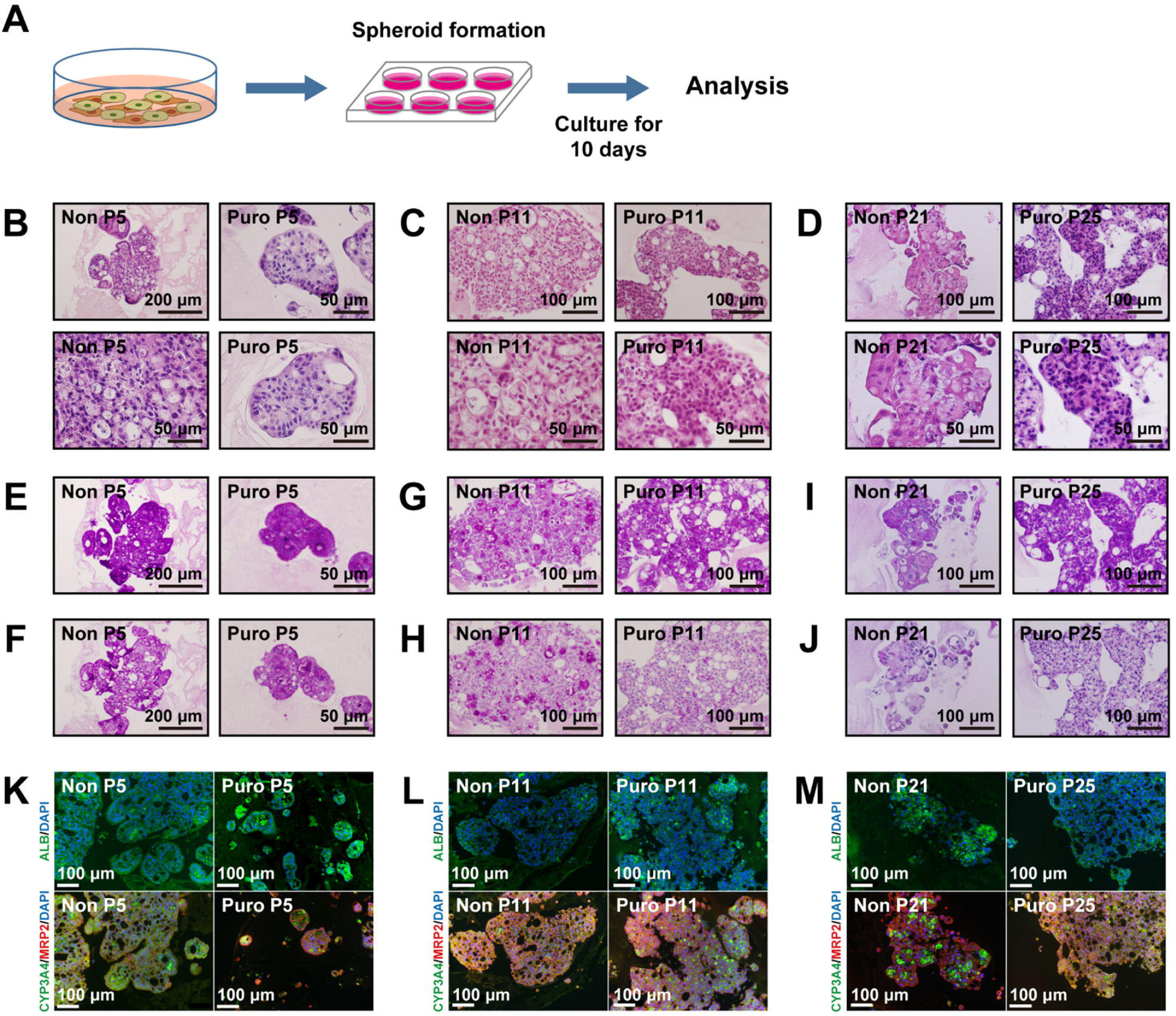
Spheroid formation induces hepatic maturation. A. Schematic of hepatic maturation protocol. Non-treated and puromycin-treated ProliHH at passage 5, 11, and 21 or 25 were cultured as spheroids for 10 days. See Figure S5A for details. B-J. Histology of spheroids generated from non-treated hepatocytes (Non) and puromycin-treated hepatocytes (Puro) at early passage (B, E, F: passage 5), middle passage (C, G, H: passage 11), and late passage (D, I, J: passage 21 or 25). B, C, D: HE stain. E, G, I: PAS stain. F, H, J: PAS stain with diastase digestion. K-M. Immunohistochemistry of spheroids generated from non-treated hepatocytes (Non) and puromycin-treated hepatocytes (Puro) at early passage (K: passage 5), middle passage (L: passage 11), and late passage (passage 21 or 25).

**Figure 7.**
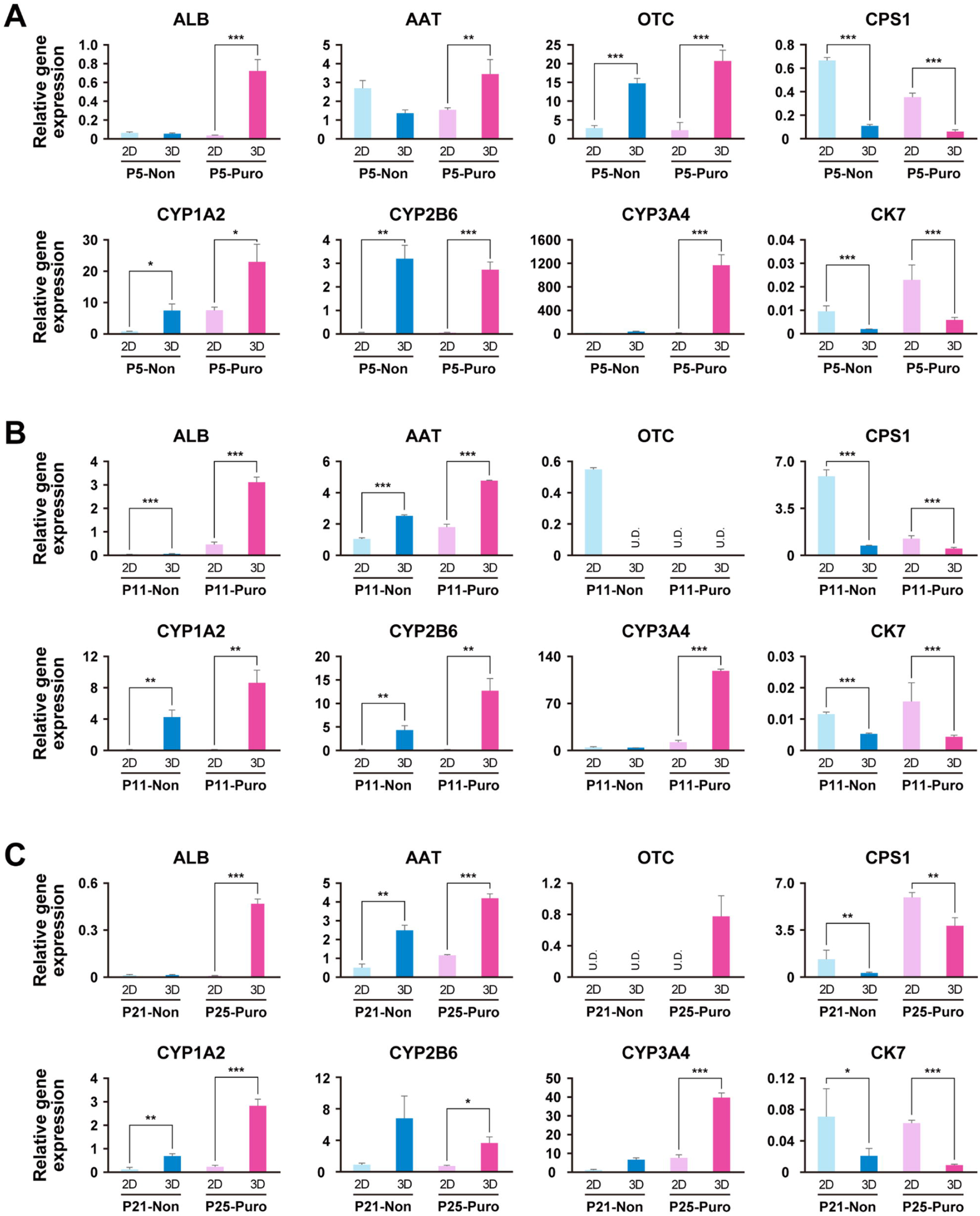
Hepatocytes restore the level of liver-associated gene expression after long-term cultivation to the levels of early stages of hepatocytic culture. Gene expression levels were analyzed by qRT-PCR. Non-treated and puromycin-treated ProliHH at passage 5 (A), 11 (B), and 21 or 25 (C) were applied to spheroid culture for 10 days (Figure S5A). The data were normalized by the housekeeping UBC gene. The expression level of each gene in passage 2 was set to 1.0. Error bars indicate the standard error. Each expression level was calculated from the results of independent (biological) triplicate experiments. The student’s T-test was performed for statistical analysis of two groups. *p < 0.05, **p < 0.01, ***p < 0.001.

## DISCUSSION

Primary human hepatocytes, especially those derived from patients with liver diseases, are useful for elucidating pathological conditions and for drug discovery. However, there is a chronic shortage of hepatocytes due to their limited proliferative capacity. In this study, the establishment of a culture system to stably grow DILI patient-derived hepatocytes enabled us to elongate hepatocyte lifespan to more than 30 population doublings with the maintenance of hepatic progenitor and progenitor-like phenotypes. Usage of Wnt3a/R-Spondin 1 in combination with feeder cells for elongation of hepatocyte lifespan is in line with a previous report that determined that WNTs are important for liver regeneration and in vitro propagation of hepatocytes and that irradiated MEF are useful for maintaining the expression and proliferative capacity of hepatic progenitors (Ito et al., 2012; Janda et al., 2017; Yanagida et al., 2013; Zhang et al., 2018).

As proof of the functionality of this culture system, our isolated hepatocytes showed resistance to puromycin, an antibiotic metabolized in the liver, while rapid cell death was induced in MEF and mesenchymal cells. This differential sensitivity to puromycin is probably due to the presence or absence of cytochrome P450 activity. Hepatic progenitors emerge after liver injury or partial hepatectomy and are responsible for liver regeneration (Li et al., 2020). In these situations where mature hepatocytes with hepatic functions decrease, hepatic progenitors survive and proliferate with drug resistance to achieve rapid and efficient liver regeneration. CYP3A4 has been implicated in the metabolism of many drugs and is an important molecular species when considering drug interactions. Computer predictions have shown that puromycin may be a substrate for CYP3A4; puromycin resistance can therefore be confirmed based on significant expression of CYP3A4. Puromycin, a commonly used antibiotic to select cells that are transduced with a puromycin-resistant gene, was useful for the selection of the proliferative biphenotypic hepatocytes and maintenance of the hepatic characteristics. Cells without the ability to metabolize the antibiotic are killed, and only functional hepatic progenitor cells or hepatocytes can metabolize the antibiotic and survive. Any antibiotic other than cell wall synthesis inhibitors may be used to select hepatocytes: Cell membrane function inhibitors, protein synthesis inhibitors, nucleic acid synthesis inhibitors, and folate synthesis inhibitors can be substituted. One type of antibiotic may be used alone, or a combination of two or more types may be used. Antibiotics that are commonly used for selecting stable transformants with exogenous genes can be used. Examples of antibiotics for selection include puromycin, blasticidin S, G418 (also known as Geneticin™), hygromycin, bleomycin, and phleomycin (also known as Zeocin™). Additionally, it is possible to isolate hepatocytes with the desired function by varying the type of additives that are specifically metabolized by hepatocytes. This functional selection method can be used like other conventional methods such as flow cytometry sorting and magnetic sorting.

The biosynthesis of ribosomes and aminoacyl-tRNAs, which are the functional binding sites of puromycin, were identified as enriched by Gene Set Enrichment Analysis. Puromycin promotes the degradation of the 80S ribosome (Aviner, 2020; Blobel and Sabatini, 1971) and the inhibition of protein synthesis by puromycin is due to competitive inhibition of aminoacyl t-RNA (Kirillov et al., 1997; Nathans and Neidle, 1963; Steiner et al., 1988). Puromycin and its derivatives bind specifically to the stop codon of full-length proteins at low concentrations, less than those at which they can compete effectively with aminoacyl-tRNA (Miyamoto-Sato et al., 2000). Therefore, in addition to the ability of hepatocytes to metabolize drugs, puromycin resistance may be attributed to the stabilization of S80 ribosome and increased aminoacyl-tRNA levels. In addition, the mitochondrial activity of puromycin-treated hepatocytes was higher than that of non-treated hepatocytes.

Furthermore, in this study, we found that the addition of puromycin maintained the proliferative capacity and functionality of the hepatocytes. In conventional culture systems, hepatocytes lose their fundamental characteristics after in vitro propagation (Katsuda et al., 2017; Wu et al., 2017; Zhang et al., 2018). Inhibition of signaling pathways that induce epithelial-mesenchymal transition, such as the TGFβ signaling pathway, is required to maintain the functionality of hepatocytes in culture (Cicchini et al., 2015; Xiang et al., 2019). Epithelial-mesenchymal transition may be involved in the reduction of functionality of the proliferating hepatocytes, and the reduction of functionality may be avoided by removing mesenchymal cells and purifying hepatocytes. Successful recovery of hepatic function upon spheroid formation indicates that the cells maintain hepatic potential during long-term propagation. Propagation of hepatocytes and subsequent maturation with spheroid formation can produce a large number of mature hepatocytes.

We herein established an efficient and stable method for the selection and expansion of hepatic progenitor-like cells by utilizing a liver-specific drug-metabolizing function. This method may lead to the widespread use of cell transplantation because of an increased availability of hepatocytes. The propagated hepatocytes could eliminate the need for natural hepatocytes. As an alternative for animal testing, propagated hepatocytes may also be amenable to scale-up to create robust platforms for drug discovery and toxicology - to test the effect of environmental pollutants, chemical compounds, and pharmaceuticals on humans. In the future, these cells may be useful for the study of human genetic diseases such as DILI and the correction of mutated genes, in combination with genome editing tools.

## Materials and Methods

### Ethical statement

All experiments handling human cells and tissues were approved by the Institutional Review Board at the National Institute of Biomedical Innovation. Informed consent was obtained from the parents of the patients. Human cells in this study were utilized in full compliance with the Ethical Guidelines for Medical and Health Research Involving Human Subjects (Ministry of Health, Labor, and Welfare, Japan; Ministry of Education, Culture, Sports, Science and Technology, Japan). Animal experiments were performed according to protocols approved by the Institutional Animal Care and Use Committee of the National Center for Child Health and Development.

### Preparation of mature hepatocytes

The liver tissues were obtained from the surplus liver of living donors from a 35-year-old woman (donor ID: 0988). Hepatocytes were isolated by the collagenase perfusion method (Enosawa, 2017; Enosawa et al., 2014; Sugahara et al., 2020). Collagenase type I (1 mg/mL, 035-17604, Fujifilm Wako Pure Chemicals, Osaka, Japan) in Hanks’ solution was used to separate hepatocytes from resected liver tissue, and liver parenchymal cells were separated by low-speed centrifugation (50 g). Cell number and viability were evaluated using the trypan blue exclusion test. The cells were frozen as cryopreserved fresh mature hepatocytes (Fresh MH) and stored in liquid nitrogen for future use.

### Preparation of hepatocytes from DILI patients

Donor identification numbers (IDs) were anonymized. The liver tissues were obtained from a 1-year-old girl with DILI (donor ID: 2064), a 7-month-old girl with DILI (donor ID: 2061), a 1-month-old girl with DILI (donor: 2062), a 7-month-old boy with DILI (donor ID: 2054), and a 9-month-old girl with DILI (donor ID: 2055). Liver tissue was shredded and incubated overnight at 37°C in DMEM (10829-018, Invitrogen; Thermo Fisher Scientific, Inc., MA, USA) containing 0.25% Collagenase Type [ (17018029, Gibco; Thermo Fisher Scientific, MA, USA) and 1% Fetal Bovine Serum (FBS: 10091148, Gibco; Thermo Fisher Scientific, MA, USA) to isolate hepatocytes. After permeabilizing the cells with a 70-μm cell strainer (352350, BD Falcon; Corning, NY, USA), hepatocytes were isolated by centrifugation at 2000 rpm for 10 min and 1000 rpm for 5 min (Low speed centrifuge LC-122, Tomy Seiko, Tokyo, Japan). The isolated cells were cultured in DMEM containing 1% penicillin-streptomycin (15140-122, Invitrogen; Thermo Fisher Scientific, MA, USA) and 10% FBS for 7-14 days, and then frozen as cryopreserved primary human hepatocytes (PHH) at 2.2 × 10^6^ cells per vial for future use (PHH2064, PHH2061, PHH2062, and PHH2055). Stem Cell Banker (CB045, Nippon Zenyaku Kogyo, Fukushima, Japan) was used as the freezing solution.

### Human hepatocyte culture and passaging

Cryopreserved PHH2064, PHH2061, and PHH2062 were used. Frozen cells were thawed and seeded onto irradiated mouse embryonic fibroblast (irrMEF) in 60 mm dishes (3010-060, IWAKI; AGC Techno Glass, Tokyo, Japan) at the seeding density of 5.0 × 10^5^ cells/cm^2^ (Passage 1). Then the cells were cultured at 37°C, 5% CO_2_ with ESTEM-HE medium (GlycoTechnica, Japan) containing Wnt3a and R-spondin 1 (Yachida et al., 2016). The medium was changed every 3 days. After 10-12 days of culture, the cells were trypsinized with 0.25% trypsin-EDTA (23315, IBL, Gunma, Japan) and plated onto 1-4 dishes in each passage.

### Preparation of feeder cells

Mouse embryonic fibroblasts (MEF) were prepared for use as nutritional support (feeder) cells. Heads, limbs, tails, and internal organs were removed from E12.5 ICR mouse fetuses (Japan CLEA, Tokyo, Japan), and the remaining torsos were then minced with a blade and seeded into culture dishes with DMEM supplemented with 10% FBS and 1% penicillin-streptomycin to allow cell growth. After 2 days of culture, the cells were passaged in a 1:4 ratio. After 5 days of culture, cells were detached with trypsin and 1/100 (v/v) of 1M HEPES buffer (15630-106, Invitrogen; Thermo Fisher Scientific, MA, USA) was added to the collected cells. Following irradiation with an X-ray apparatus (dose: 30 Gy, MBR-1520 R-3, Hitachi, Tokyo, Japan), the cells were frozen using TC protector (TCP-001DS, Pharma Biomedical, Osaka, Japan).

### Puromycin selection

For hepatocyte selection, puromycin (final concentration: 1 μg/mL or 2 μg/mL, 160-23151, FUJIFILM Fujifilm Wako Pure Chemicals, Osaka, Japan) was exposed to proliferating hepatocytes for 3 days. After exposure to puromycin, cells were washed with PBS (14190-250, Invitrogen; Thermo Fisher Scientific, MA, USA) and cultured in fresh ESTEM-HE medium (GlycoTechnica, Japan) for at least 1 day. When the cells reached 90% confluence, the cells were passaged by 0.25% trypsin-EDTA treatment.

### human iPSC culture

Induced pluripotent stem cells (iPSCs) were generated from fibroblasts of the donor 2064 (iPSC-O) and the donor 2054 (iPSC-K) by the introduction of the Sendai virus carrying the 4 Yamanaka factors (Takahashi and Yamanaka, 2006). The iPSCs were cultured on irrMEF with medium for human iPSCs : KnockoutTM-Dulbecco’s modified Eagle’s medium (KO-DMEM: 10829-018, Life Technologies; Thermo Fisher Scientific, MA, USA) supplemented with 20% KnockoutTM-Serum Replacement (KO-SR: 10828-028, Gibco; Thermo Fisher Scientific, MA, USA), 2 mM Glutamax-I (35050-079, Gibco; Thermo Fisher Scientific, MA, USA), 0.1 mM non-essential amino acids (NEAA)(11140-076, Gibco; Thermo Fisher Scientific, MA, USA), 1% penicillin-streptomycin (Invitrogen), 0.055 mM 2-Mercaptoethanol (21985-023, Invitrogen; Thermo Fisher Scientific, MA, USA) and recombinant human full-length bFGF (PHG0261, Gibco; Thermo Fisher Scientific, MA, USA) at 10 ng/ml.

### Hepatic differentiation of human iPSCs

Differentiation to hepatocyte-like cells from iPSC-K (HLC-KI): To generate embryoid bodies (EBs), iPSC-K (1 × 10^4^ cells/well) were dissociated into single cells with Accutase (A1110501, Gibco; Thermo Fisher Scientific, MA, USA) after exposure to ROCK inhibitor, Y-27632 (A11105-01, Fujifilm Wako Pure Chemicals, Osaka, Japan), and cultivated in 96-well plates (174925, Thermo Fisher Scientific, MA, USA) in EB medium [75% KO-DMEM, 20% KO-SR, 2 mM GlutaMAX-I, 0.1 mM NEAA, 1% penicillin-streptomycin, and 50 μg/mL L-ascorbic acid 2-phosphate (A8960, Sigma-Aldrich, MO, USA)] for 10 days. The EBs were transferred to the 24-well plates coated with collagen type I and cultivated in XF32 medium [85% Knockout DMEM, 15% Knockout Serum Replacement XF CTS (XF-KSR: 12618013, Gibco; Thermo Fisher Scientific, MA, USA), 2 mM GlutaMAX-I, 0.1 mM NEAA, 1% penicillin-streptomycin, 50 μg/mL L-ascorbic acid 2-phosphate, 10 ng/mL heregulin-1β, 200 ng/mL recombinant human IGF-1 (LONG R3-IGF-1: 85580C, Sigma-Aldrich, MO, USA), and 20 ng/mL human bFGF] for 35 days. For iPSC-O differentiation to hepatocyte-like cells (HLC-O): Hepatic differentiation of iPSC-O was performed by Cellartis Hepatocyte Differentiation Kit (Y30050, Takara Bio, Shiga, Japan) according to the manufacturer’s instructions. In this study, cells were used after 30 days of differentiation induction.

### Calculation of population doublings

Cells were harvested at sub-confluency and the total number of cells in each well was determined using a cell counter. Population doubling was used as the measure of cell growth. PD was calculated from the formula PD=log_2_(A/B), where A is the number of harvested cells and B is the number of plated cells (Akutsu et al., 2015).

### Histology and Periodic Acid Schiff (PAS) staining

Samples were coagulated in iPGell (PG20-1, GenoStaff, Tokyo, Japan) following the manufacturer’s instructions and fixed in 4% paraformaldehyde at 4°C overnight. Fixed samples were embedded in a paraffin block to prepare cell sections. For Hematoxylin Eosin (HE) staining, the deparaffinized sections were treated with a hematoxylin solution (Mutoh Chemical, Tokyo, Japan) for 5 min at room temperature and washed with dilute ammonia. After washing with 95% ethanol, dehydration was performed with 150 mL of eosin in 95% ethanol solution and permeabilized in xylene. For PAS staining, the deparaffinized sections were reacted with 0.5% periodate solution (86171, Mutoh Chemical, Tokyo, Japan) for 10 min at room temperature, rinsed with water for 7 min. After reacting with Schiff’s reagent (40922, Mutoh Chemical, Tokyo, Japan) for 5-15 min, the sections were washed with sulfurous acid water. Coloration was achieved by reaction with Meyer hematoxylin solution (30002, Mutoh Chemical, Tokyo, Japan) for 2 min at room temperature and then rinsing with water for 10 min.

### Senescence-associated β-galactosidase staining

For senescence-associated β-galactosidase staining, cells were fixed in 4% paraformaldehyde for 10 min at room temperature. Fixed cells were stained with the Cellular Senescence Detection Kit (CBA-230, Cell Biolabs, CA, USA) following the manufacturer’s instructions.

### Karyotypic analysis

Karyotypic analysis was contracted out to Nihon Gene Research Laboratories (Sendai, Japan). Metaphase spreads were prepared from cells treated with 100 ng/mL of Colcemid (Karyo Max, Gibco; Thermo Fisher Scientific, MA, USA) for 6 h. The cells were fixed with methanol: glacial acetic acid (2:5) three times and placed onto glass slides. Giemsa banding was applied to metaphase chromosomes. A minimum of 10 metaphase spreads was analyzed for each sample and karyotyped using a chromosome imaging analyzer system (Applied Spectral Imaging, CA, USA).

### qRT-PCR

Total RNA was prepared using ISOGEN (311-02501, Nippon Gene, Tokyo, Japan) and RNeasy Micro Kit (74004, Qiagen, Hilden, Germany). The RNA was reverse transcribed to cDNA using Superscript ◻ Reverse Transcriptase (18080-085, Invitrogen; Thermo Fisher Scientific, MA, USA) with ProFlex PCR System (Applied Biosystems, MA, USA). Quantitative RT-PCR was performed on QuantStudio 12K Flex (Applied Biosystems, MA, USA) using a Platinum SYBR Green qPCR SuperMix-UDG (11733046, Invitrogen; Thermo Fisher Scientific, MA, USA). Expression levels were normalized with the reference gene, *UBC*. The primer sequences are shown in Table 1.

**Table 1.**
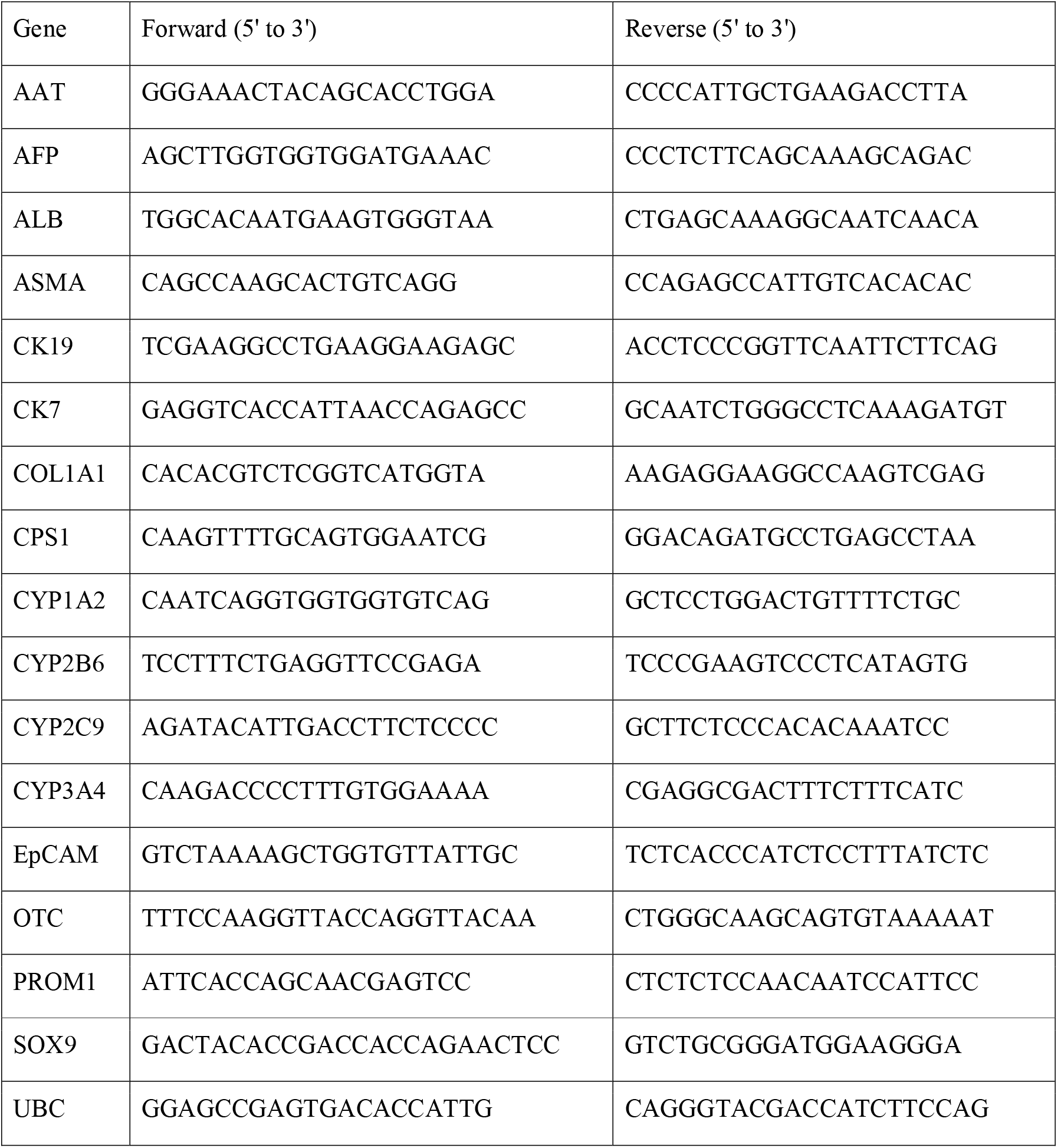
Primers used for qRT-PCR.

### Immunofluorescence staining

Cells were fixed with 4% paraformaldehyde in PBS for 10 min at room temperature. After washing with PBS, cells were permeabilized with 0.1% Triton X in PBS for 10 min, pre-incubated with Protein Block Serum-Free (X0909, Dako, Jena, Germany) for 30 min at room temperature and then exposed to primary antibodies overnight at 4°C. Cells were washed with PBS and incubated with diluted secondary antibodies for 30 min at room temperature. Nuclei were stained with 4′,6-diamidino-2-phenylindole, dihydrochloride (DAPI, 40043, Biotium, CA, America). For immunofluorescence staining of cut paraffin sections, sections were deparaffinized for 30 min before staining. Then sections were rinsed with distilled water for 3 min, and antigen retrieval was performed for 20 min using heated Histophine diluted 10 times with distilled water. After standing at room temperature for 20 min, sections were washed three times with PBS. Endogenous peroxidase removal was performed using 3% hydrogen peroxide water diluted 10 times with methanol for 5 min. After washing three times with PBS, sections were incubated with primary antibodies overnight at 4°C. After washing three times with PBS, sections were incubated with diluted secondary antibodies for 30 min at room temperature. Nuclei were stained with DAPI. The antibodies listed in Table 2 and 3 were diluted according to the tables in PBS containing 1% BSA (126575, Calbiochem).

**Table 2.**
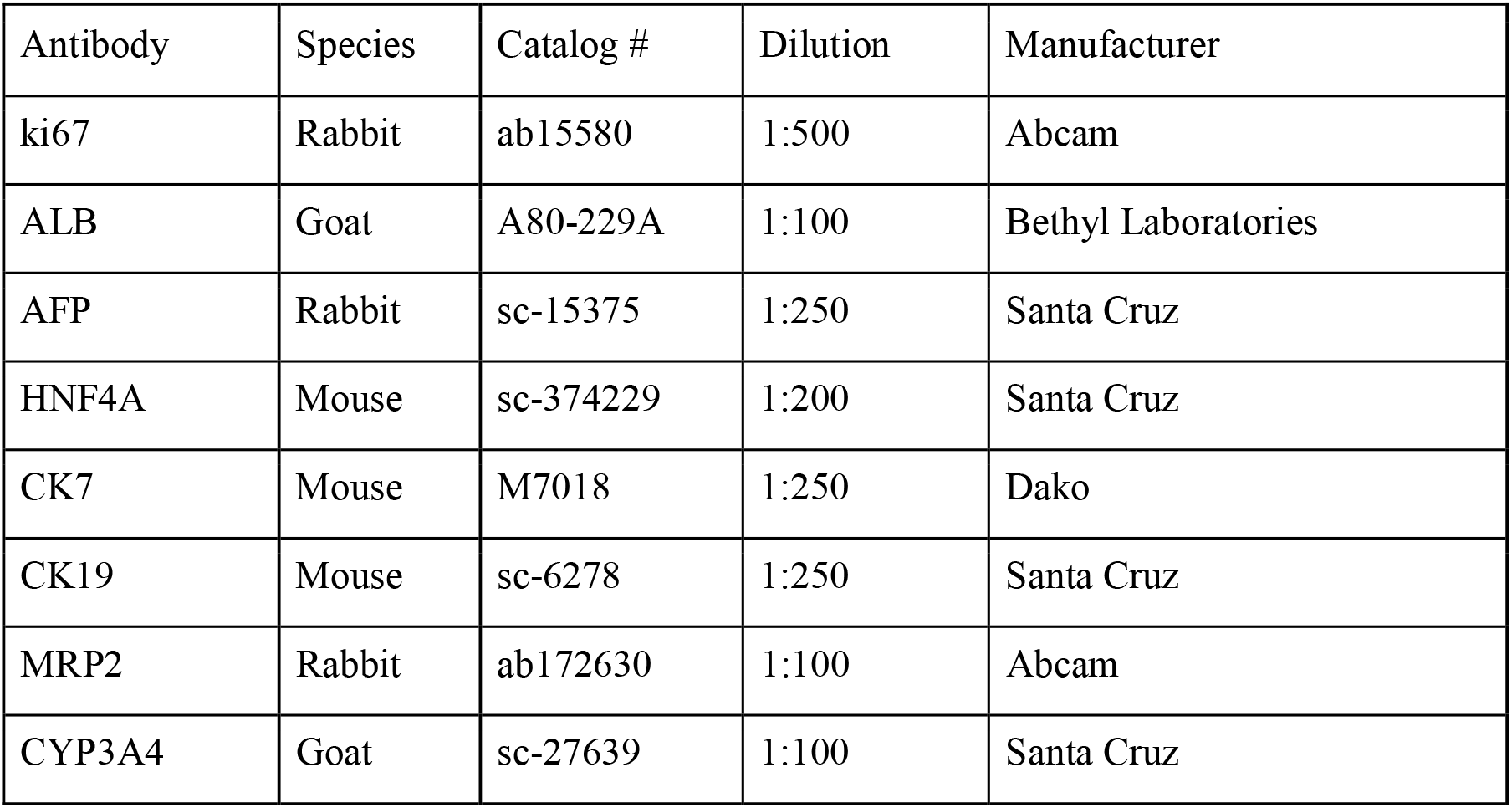
Antibodies used for immunohistochemistry.

**Table 3.** secondary antibodies used for immunohistochemistry.

| Antibody | Manufacturer | Catalog # |
| --- | --- | --- |
| Alexa Fluor 488 rabbit anti-goat IgG | Invitrogen | A11078 |
| Alexa Fluor 488 goat anti-rabbit IgG | Invitrogen | A11008 |
| Alexa Fluor 488 goat anti-mouse IgG | Invitrogen | A21121 |
| Alexa Fluor 546 goat anti-mouse IgG1 | Invitrogen | A21123 |
| Alexa Fluor 568 goat anti-mouse IgG2a | Invitrogen | A21134 |

### Cytochrome P450 induction

To examine the induction of cytochrome P450 (CYP) enzymes, cells were cultured in 12-well plates (353043, BD Falcon; Corning, NY, USA) using irrMEF and ESTEM-HE medium. When the cells reached 80-90% confluence, the following drugs were added to the cells: 20 μM rifampicin (solvent: DMSO, 189-01001, Fujifilm Wako Pure Chemicals, Osaka, Japan) with an induction period of 2 days, 50 μM omeprazole (solvent: DMSO, 158-03491, Fujifilm Wako Pure Chemicals, Osaka, Japan) with an induction period of 1 day, and 500 μM phenobarbital (solvent: DMSO, 162-11602, Fujifilm Wako Pure Chemicals, Osaka, Japan) with an induction period of 2 days.

### Measurement of CYP3A4 activity

For the measurement of CYP3A4 activity, cells were cultured in irrMEF and ESTEM-HE until 90% confluence. CYP3A4 activity was analyzed by P450-Glo CYP3A4 Assay and Screening System (V9001, Promega, WI, USA) following the manufacturer’s instructions. HepG2 cells (HEPG2-500, Cellular Engineering Technologies, IA, USA) cultured in HepG2 Hepatocellular Carcinoma Expansion Media ( HEPG2.E.MEDIA-450, Cellular Engineering Technologies, IA, USA) were used as a positive control.

### Hepatic maturation by three-dimensional (3D) culture

For hepatic maturation, the cells were detached with 0.25% trypsin/EDTA, transferred to 6-well plates (3471, Corning, NY, USA) at a density of 5 × 10^5^ cells/well, and then cultivated for 10 days to form spheroids. Fresh medium was added at day 5 and analyses were performed at day 10.

### Microarray analysis

Total RNA was isolated using miRNeasy mini kit (217004, Qiagen, Hilden, Germany). RNA samples were labeled and hybridized to a SurePrint G3 Human GEO microarray 8 × 60K Ver 3.0 (Agilent, CA, USA), and the raw data were normalized using the 75-percentile shift. For gene expression analysis, a one-way ANOVA was performed to identify differentially expressed genes (DEGs). Fold-change numbers were calculated for each analysis (p-value < 0.05, fold-change>1.5). Unsupervised clustering was performed with sorted or whole DEGs using the R package. Gene expression profiles of mature hepatocytes were analyzed using human liver total RNA (636531, Clontech: Takara Bio, Shiga, Japan). Expression profiles of bile duct epithelial cells were obtained from Gene Expression Omnibus (GSM4454532, GSM4454533, GSM4454534). Principal component analysis was performed with whole genes using the R package. Functional enrichment analysis including Over-Representation Analysis (ORA) and Gene Set Enrichment Analysis was performed by using WebGestalt (http://www.webgestalt.org/).

### Statistical analysis

The numbers of biological and technical replicates are shown in the figure legends. All data are presented as mean ± SD (technical triplicate) or mean ± SE (biological triplicate). For most statistical evaluations in this study, an unpaired Student’s t-test was used to calculate statistical probabilities. p-values were calculated by two-tailed t-test.

For gene expression analysis, one-way ANOVA was performed to identify differentially expressed genes (DEGs), and the fold-change number was calculated for each analysis (p-value < 0.05, fold-change > 1.5).

## Acknowledgments

We would like to express our sincere thanks to K. Miyado for fruitful discussion, to M. Ichinose and K. Tatsumi for providing expert technical assistance, to C. Ketcham for English editing and proofreading, and E. Suzuki and K. Saito for secretarial work.

## Author Contribution Statement

AU and SA designed the experiments. SA, NS, SH, and KI performed the experiments. SA, NS, SM, and AU analyzed data. MT, TKim, Mku, AN, MKa, HN, AK, TKiy, and YM contributed to the reagents, tissues, and analysis tools. SA, MT, TKim, Mku, HN, AK, TKiy, YM, and AU discussed the data and manuscript. SA and AU wrote this manuscript.

## Supplemental Figure Legends

**Figure S1. Details of the morphology and gene expression of proliferating hepatocytes, related to Figures 1 and 2**.

A. Phase-contrast photomicrographs of primary human hepatocytes on irrMEF for 7 days. Two colonies are shown at low and high magnifications.

B. Phase-contrast photomicrographs of ProliHH from passage 5.

C. Gene expression by qRT-PCR in primary human hepatocytes (PHH) at passage 1 and 2 (Hep2064 P1 and P2). The data were normalized with the housekeeping UBC gene. Each relative value was calculated with respect to HepG2. Error bars indicate the standard deviation (n=3). “HepG2 (human hepatoma cell)”, “Ad_Liver (human adult normal liver pools of five donors purchased from BioChain, R1234149-P)”, and “Fresh_MH” (hepatocytes isolated from the adult human liver in our laboratory) are used for comparison.

D. Gene expression by qRT-PCR in ProliHH from passage 2 to 21. The data were normalized with the housekeeping UBC gene. The expression level of each gene in passage 2 was set to 1.0. Error bars indicate the standard deviation (n=3).

**Figure S2. Evaluation of the effect of puromycin on mouse fetal fibroblasts (MEF) and hepatocytes derived from other DILI patients, related Figure 3**.

A. Phase-contrast photomicrographs of MEF with exposure of puromycin (Puro: 0, 1, 2, 10, 50, and 100 μg/mL) for 3 days. Puromycin was added at 100% confluence.

B. Phase-contrast photomicrographs of puromycin-treated ProliHH at passage 5.

C-E. Phase-contrast photomicrographs of PHH (#2062) 3 days after exposure to 2 μg/mL puromycin. Puromycin was added when the cells reached confluence (Day 0) and removed 3 days after the addition (Day 3). The Day 4 image shows the cells 24 hours after puromycin removal (D).

F. Gene expression by qRT-PCR in puromycin-treated hepatocytes (#2061 and #2062). The data were normalized by the housekeeping UBC gene. From left to right: non-treated hepatocytes (Hep2061), puromycin-treated hepatocytes (Hep2061+puro), non-treated hepatocytes (Hep2062), and puromycin-treated hepatocytes (Hep2062+puro). Cells were treated with 2 μg/mL puromycin for 3 days. Each relative value was calculated with respect to non-treated cells. Error bars indicate the standard deviation (n=3). The student’s T-test was performed for statistical analysis of two groups. *p < 0.05, **p < 0.01, ***p < 0.001.

**Figure S3. Details of gene expression analysis and karyotyping of puromycin-treated proliferating hepatocytes and hepatocyte selection with a low concentration (1 μg/mL) of puromycin, related to Figures 3 and 4**.

A. Gene expression by qRT-PCR in puromycin-treated ProliHH from passage 4 to 25. The data were normalized with the housekeeping UBC gene. The expression level of each gene at passage 2 was set to 1.0. Error bars indicate the standard deviation (n=3).

B. Karyotypes of puromycin-treated ProliHH at passage 24. About 80% of the hepatocytes were 46XX and no major abnormalities were found.

C. Growth curves of hepatocytes treated with 1μg/mL puromycin (n=2, shown as green and yellow dots). Proliferative capacity was analyzed at each passage. Cells were passaged in the ratio of 1:4 for each passage. The numbers of cells were calculated as an average of 50 counts. “Population doubling” indicates the cumulative number of divisions of the cell population.

D. Phase-contrast photomicrographs of puromycin-treated ProliHH from passages 5, 7, 13, 21, and 25.

E. Microscopic view of ProliHH at passages 5, 13, and 25. HE stain.

F. Immunocytochemical analysis of puromycin-treated ProliHH with an antibody to Ki67 (cell proliferation marker).

G. A senescence-associated beta-galactosidase stain of puromycin-treated ProliHH at the indicated passages. The number of β-galactosidase-positive senescent cells increased at passage 25.

H. Karyotypes of puromycin-treated ProliHH at passage 24.

**Figure S4. Long-term analysis of hepatocytes selected with a low concentration (1 μg/mL) of puromycin, related to Figures 3 and 4**.

A. Gene expression by qRT-PCR in ProliHH at passage 4, 11, and 17. The data were normalized by the housekeeping gene UBC. The expression level of each gene at passage 2 was set to 1.0. Error bars indicate the standard deviation (n=3). The student’s T-test was performed for statistical analysis of two groups. *p < 0.05, **p < 0.01, ***p < 0.001.

B. Gene expression by qRT-PCR in puromycin-treated ProliHH from passage 4 to 25. Data were normalized with the housekeeping gene UBC. The expression level of each gene at passage 2 was set to 1.0. Error bars indicate the standard deviation (n=3).

C-E. Immunocytochemical analysis of puromycin-treated ProliHH at a low concentration ( (1μg/mL) with antibodies to ALB (C), HNF4A (D), and bile duct marker CK7 (E) at early and late passages (passage 5 and 25, respectively). Yellow arrows indicate binuclear cells (E).

F-G. Glycogen storage in puromycin-treated hepatocytes (passage 5, 13, and 25) by PAS stain with (F) and without (G) diastase digestion.

H-J. Expression of the genes for CYP1A2 (H), CYP2B6 (I), and CYP3A4 (J) in puromycin-treated hepatocytes after exposure to omeprazole (H), phenobarbital (I), or rifampicin (J). The expression level of each gene without any treatment (DMSO) was set to 1.0. Each expression level was calculated from the results of independent (biological) triplicate experiments. Error bars indicate the standard error (n=3). The student’s T-test was performed for statistical analysis of two groups. *p < 0.05.

K. CYP3A4 activity of hepatocytes (passage 5, 11, and 25). The CYP3A4 activity of HepG2 cells was set to 1.0. Error bars represent the standard errors (n=3). Each expression level was calculated from the results of independent (biological) triplicate experiments. Error bars indicate the standard error (n=3). *p < 0.05, NS: p > 0.05.

**Figure S5. Detailed expression analysis and comprehensive analysis of hepatic progenitor cell markers in proliferating hepatocytes, related to Figure 5**.

A, B. Expression of the genes for hepatic progenitor cell markers by qRT-PCR in non-treated (A) and puromycin-treated (B) ProliHH. Data were normalized with the housekeeping gene UBC. The expression level of each gene in HepG2 cells was set to 1.0, as a positive control. Error bars indicate the standard deviation (n=3).

C. Immunocytochemical analysis of non-treated and puromycin-treated ProliHH with antibodies to AFP and CK19 at the early passage (passage 5).

D-F. Gene sets enriched in either (FDR < 0.05, P < 0.05) compared to non-treated and puromycin-treated ProliHH. Gene sets for the TCA cycle and pentose phosphate circuit (D), and epithelial-mesenchymal transition (EMT) (E) were identified. “non”: non-treated ProliHH, “+puro”: puromycin-treated ProliHH.

**Figure S6. Hepatocytes treated with low concentrations (1 μg/mL) of puromycin restore the expression level of liver-related genes after long-term culture to the level of the initial stage of hepatocyte culture, related to Figure 6 and 7**.

A. Hepatic maturation protocol. Hepatocytes at early passage (Passage 5), middle passage (Passage 11), and late passage (Passage 25) at 70% confluence were exposed to 1 or 2 μg/mL puromycin for 3 days. Hepatocytes were then seeded onto low adhesion plates for spheroid formation one day after removal of puromycin. The cells were cultured at a form of spheroid for 10 days and subjected to further analysis.

B-J. Histology of spheroids generated from ProliHH at each passage (B, E, F: Early passage (Passage 5); C, G, H: middle passage (Passage 11); D, I, J: late passage (Passage 25)). “+puro (x1)”: Spheroids generated from ProliHH treated with 1 μg/ml puromycin. B to D: HE stain; E, G, I: PAS stain; F, H, J: PAS stain used in combination with diastase, an enzyme that breaks down glycogen.

K-M. Immunohistochemistry of spheroids generated from puromycin-treated ProliHH at each passage (K: Early passage (passage 5); L: middle passage (passage 11); M: late passage (passage 25)) with antibodies to albumin (ALB), cytochrome p450 3A4 (CYP3A4), and multidrug resistance-associated protein 2 (MRP2). “+puro (x1)”: Spheroids generated from ProliHH treated with 1 μg/ml puromycin.

**Figure S7. Hepatocytes treated with low concentrations (1 μg/mL) of puromycin restore the expression level of liver-related genes after long-term culture to the level of the initial stage of hepatocyte culture, related to Figure 6 and 7**.

Gene expression levels were analyzed by qRT-PCR. Puromycin-treated ProliHH were subjected to spheroid culture conditions for 10 days (Figure S6A). RNAs were isolated from spheroids generated from puromycin-treated ProliHH at passage 5 (A), passage 11 (B), and passage 25 (C). The data were normalized by the housekeeping gene UBC. The expression level of each gene in ProliHH without any treatment at passage 2 was set to 1.0. “U.D.”: undetectable. Each expression level was calculated from the results of independent (biological) triplicate experiments. Error bars indicate the standard error. The student’s T-test was performed for statistical analysis of two groups. *p < 0.05, **p < 0.01, ***p < 0.001.

