## Supplementary figures and images for "Drug metabolic activity is a critical cell-intrinsic determinant for selection of hepatocytes during long-term culture"

### Supplemental Figure 1

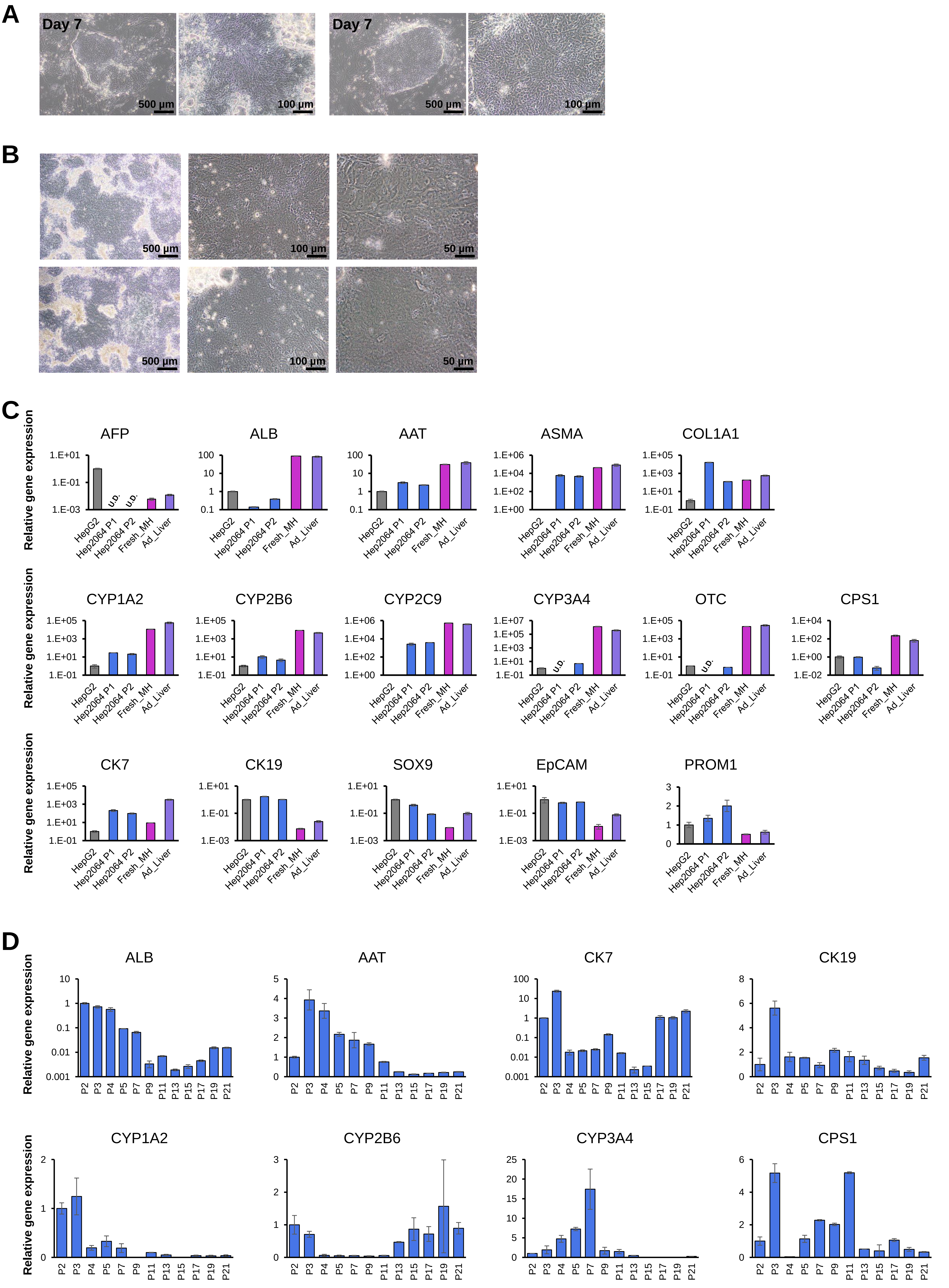

### Supplemental Figure 2

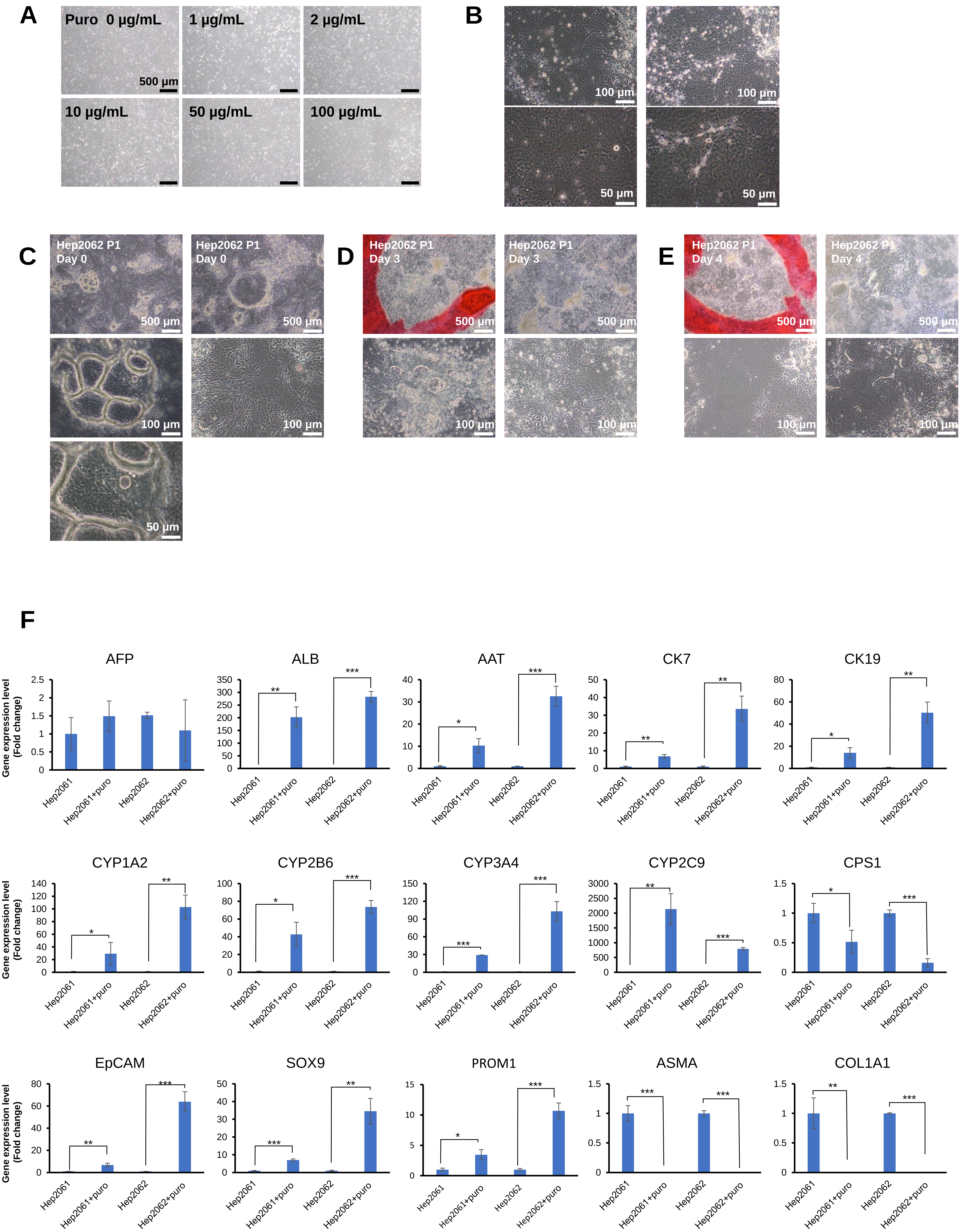

### Supplemental Figure 4

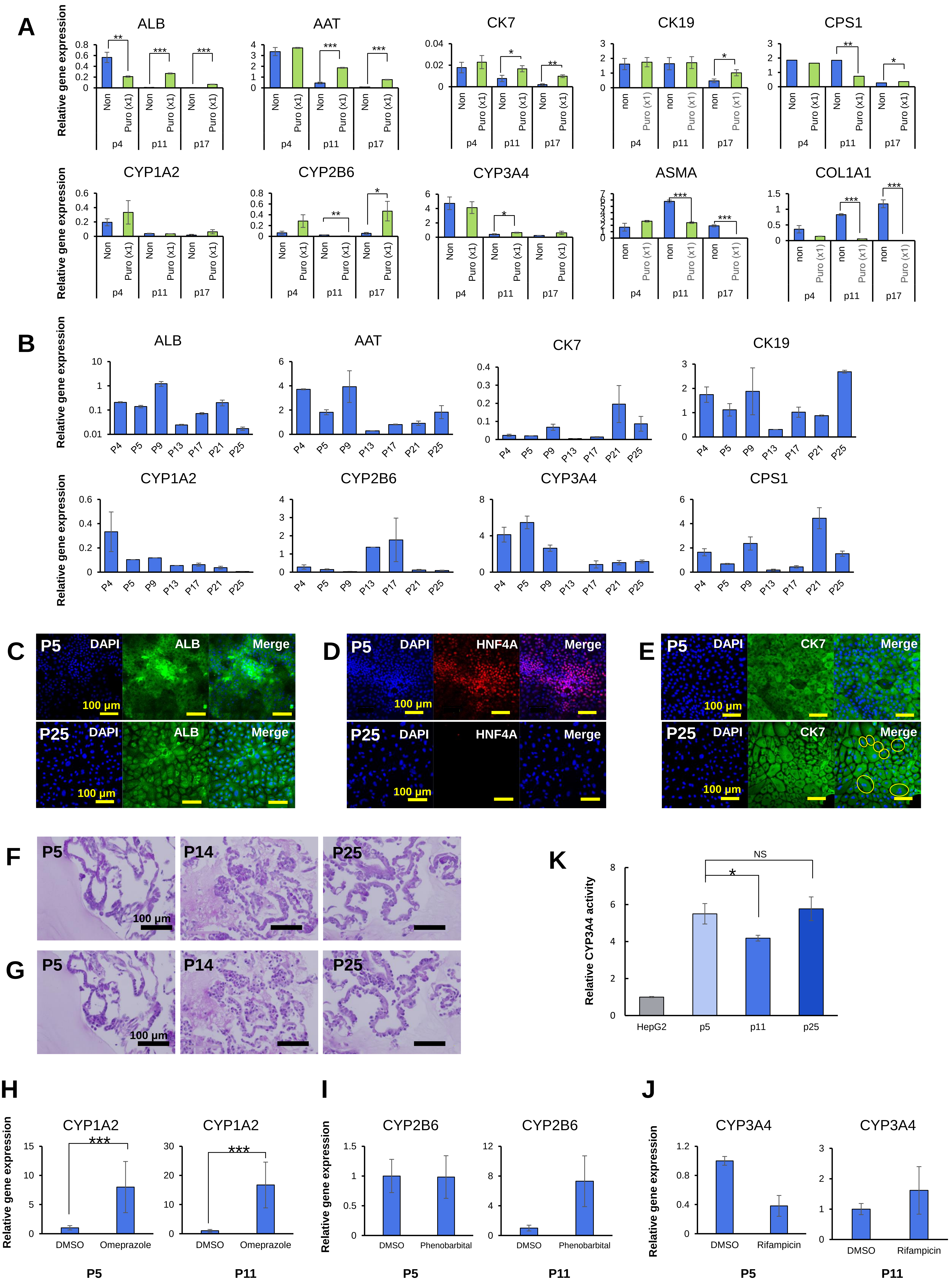

### Supplemental Figure 5

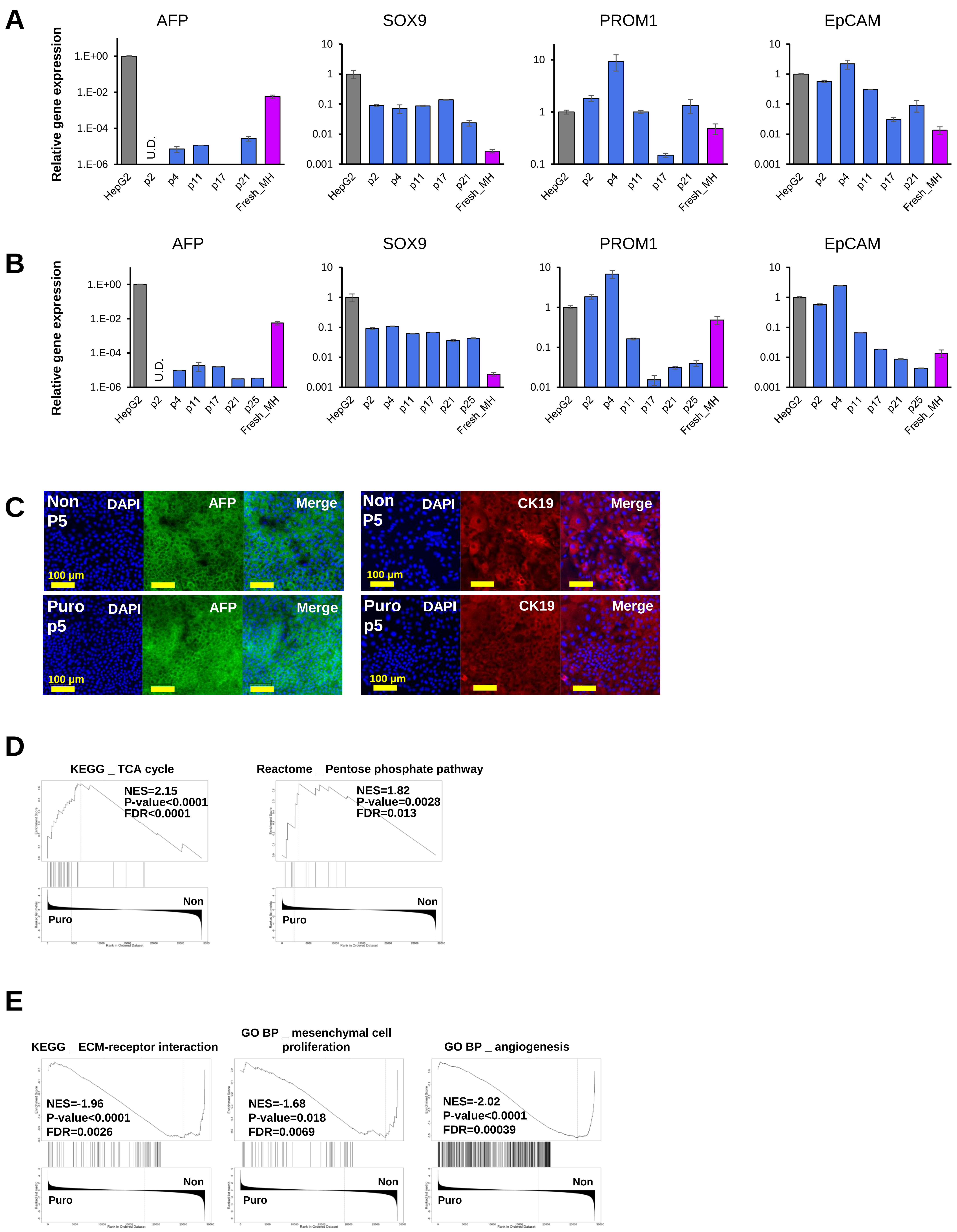

### Supplemental Figure 6

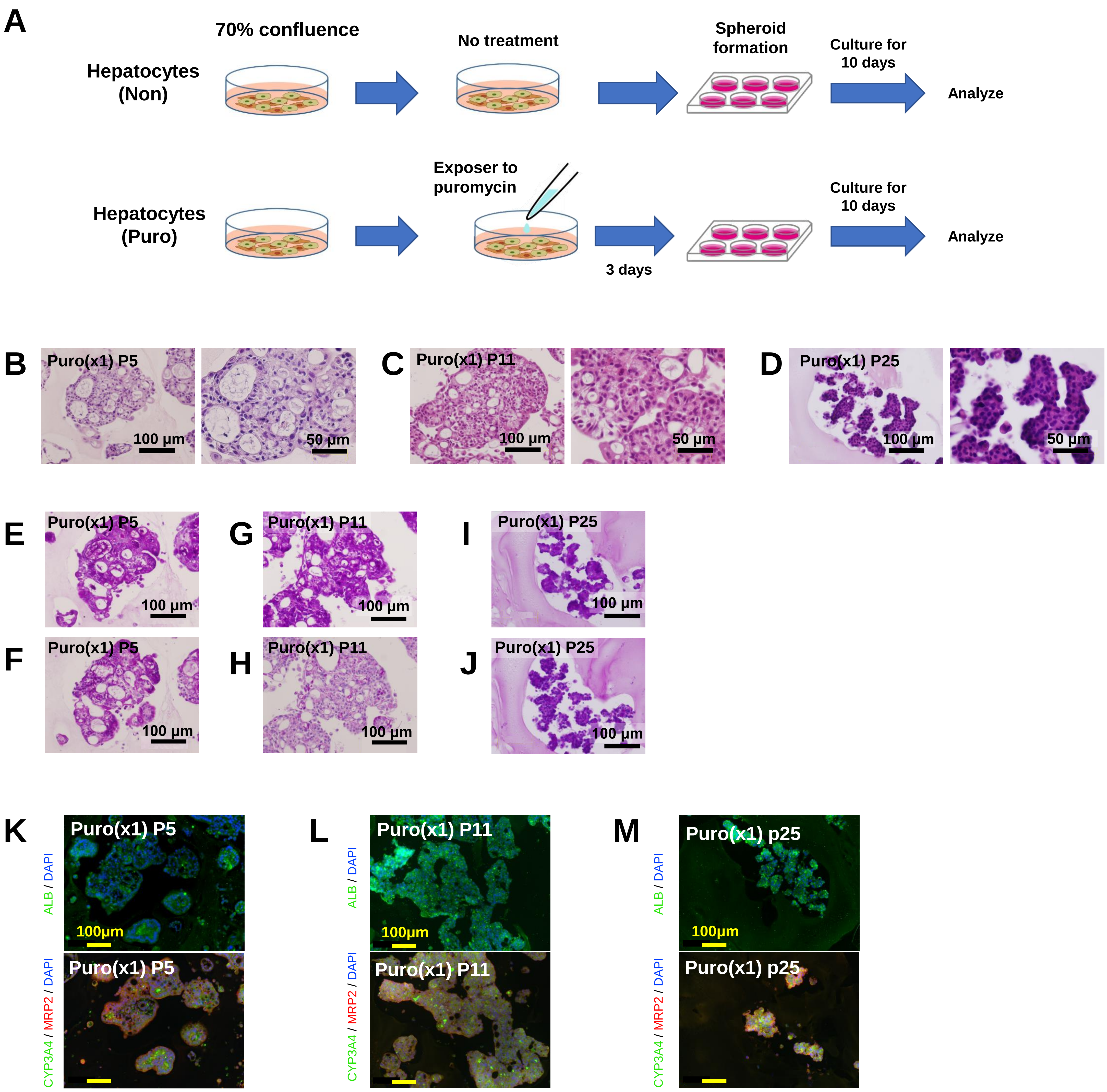

### Supplemental Figure 7

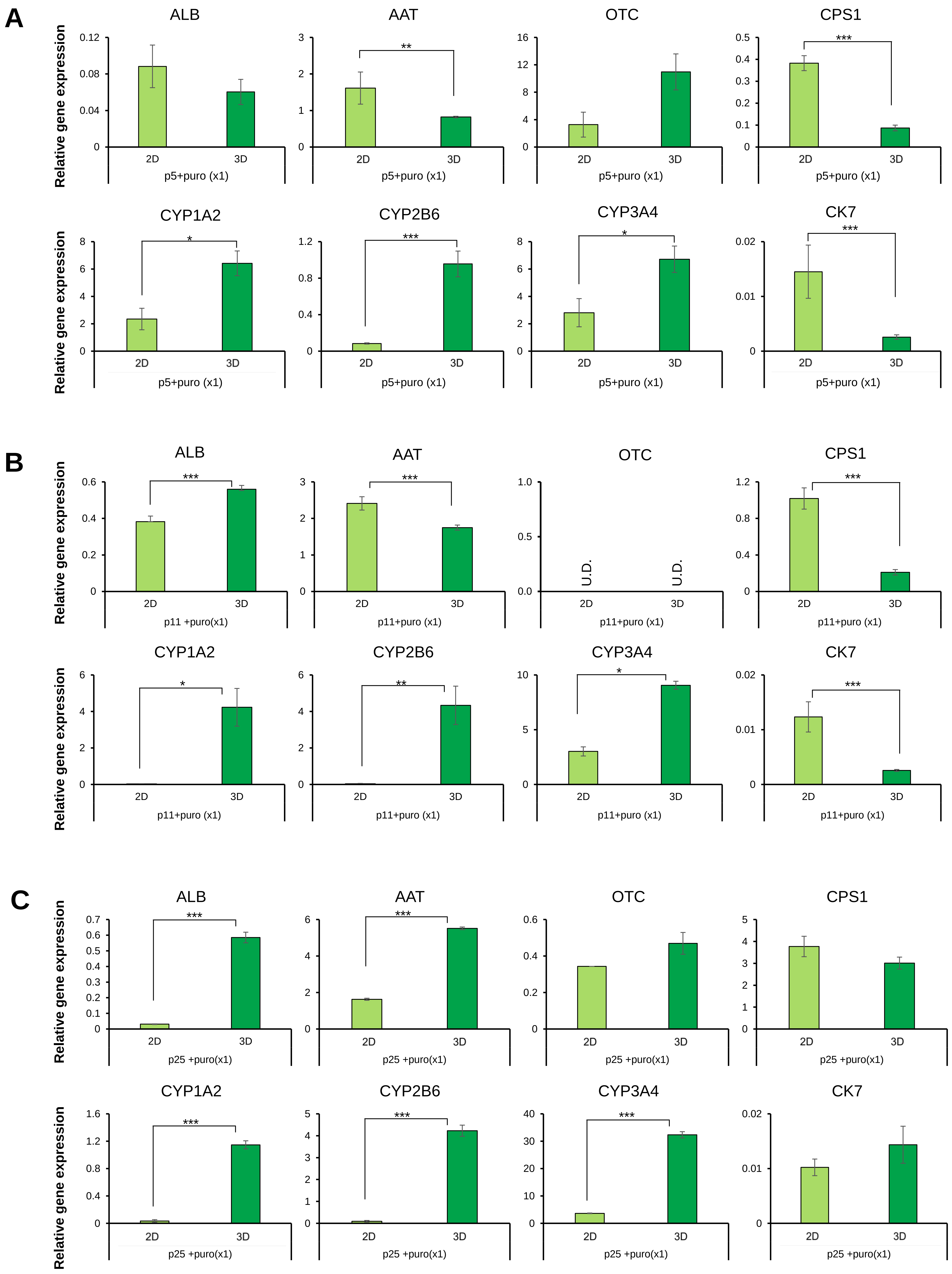
