## Supplemental Figure 3 for "Drug metabolic activity is a critical cell-intrinsic determinant for selection of hepatocytes during long-term culture"

A

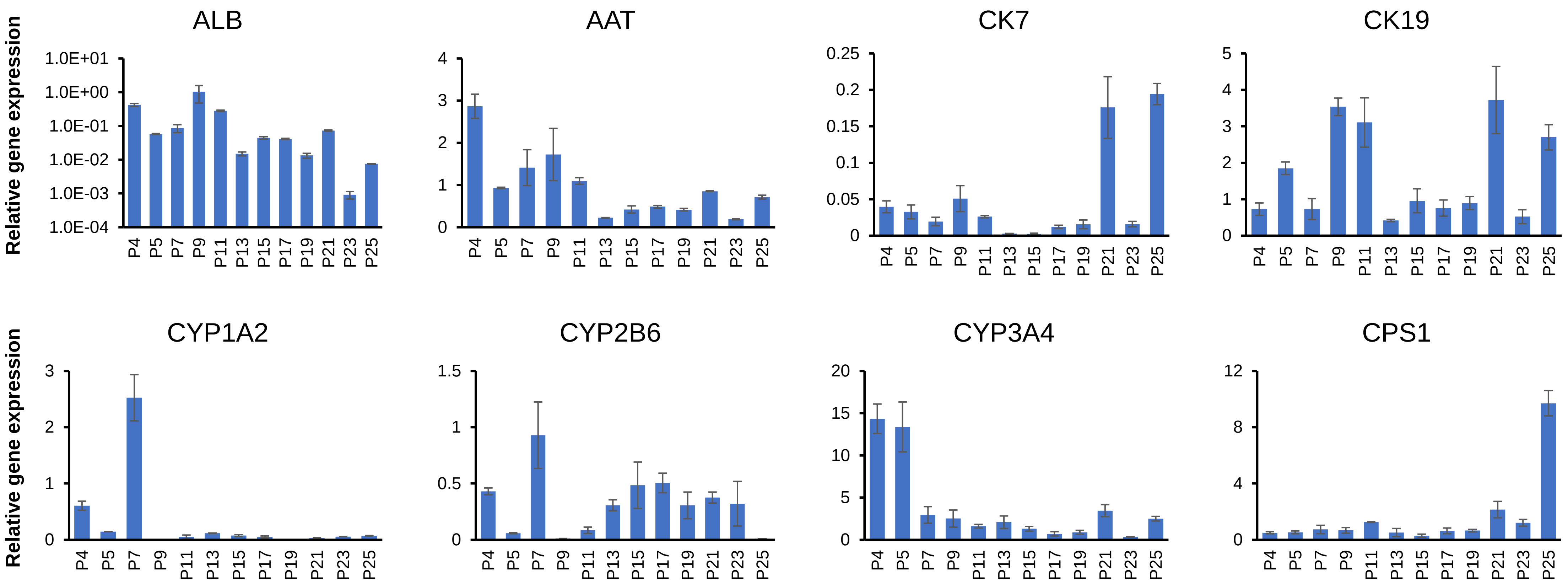


B

| Karyotype | Cell number |
|---|---|
| 46,XX | 14 |
| 45,XX,dic(14;17)(p11.2;p13) | 3 |
| 45,XX,dic(1314)(q34;q32) | 1 |
| 45,XX,dic(5;14)(p15.3-p11.2),add(17)(p13) | 1 |
| 45,XX,add(14)(q32),-18,-18,+mar | 1 |
| | 20 |

| Chromosome number | Cell number |
|---|---|
| 46 | 42 |
| 45 | 8 |
| | 50 |

C

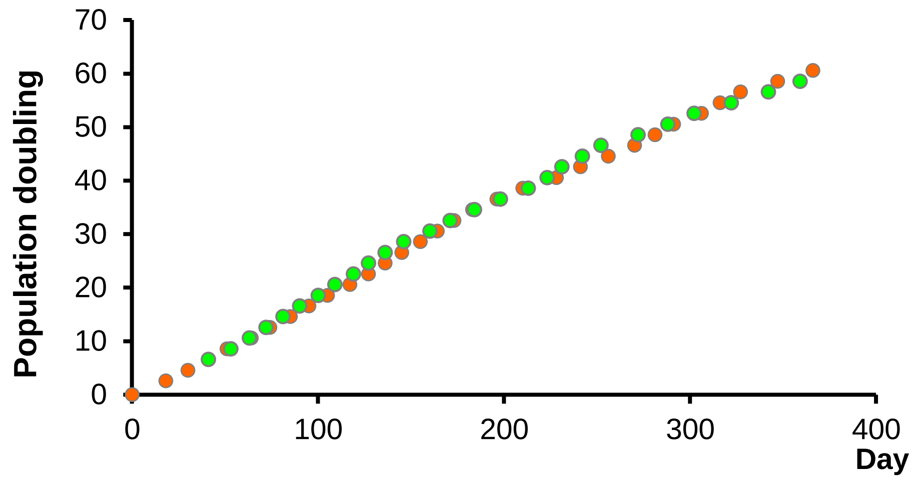


D

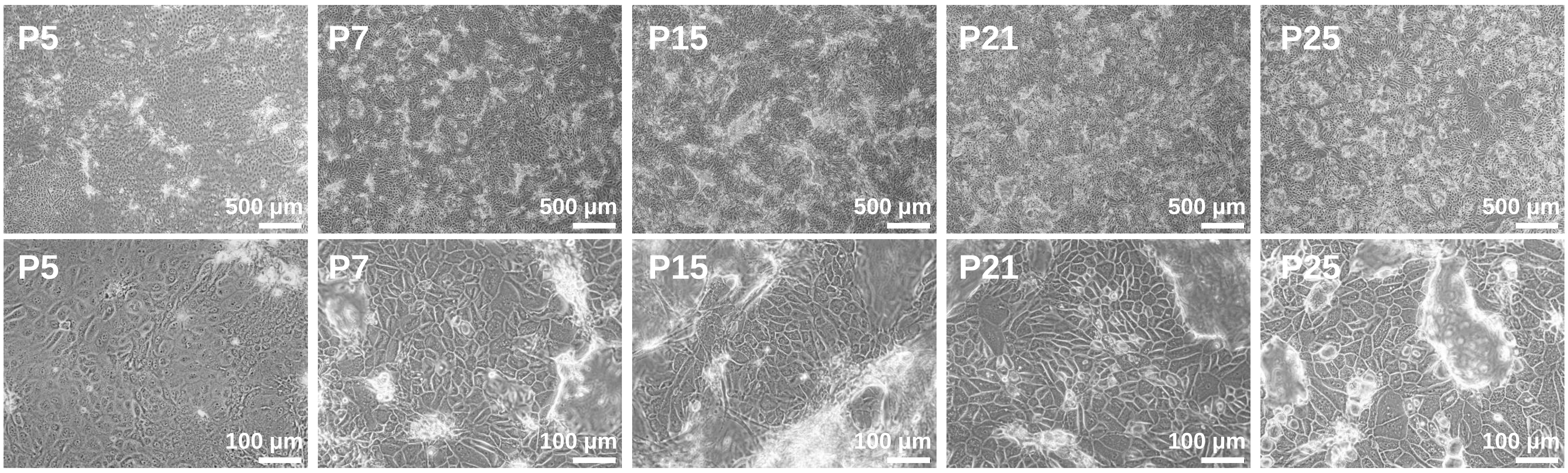


E

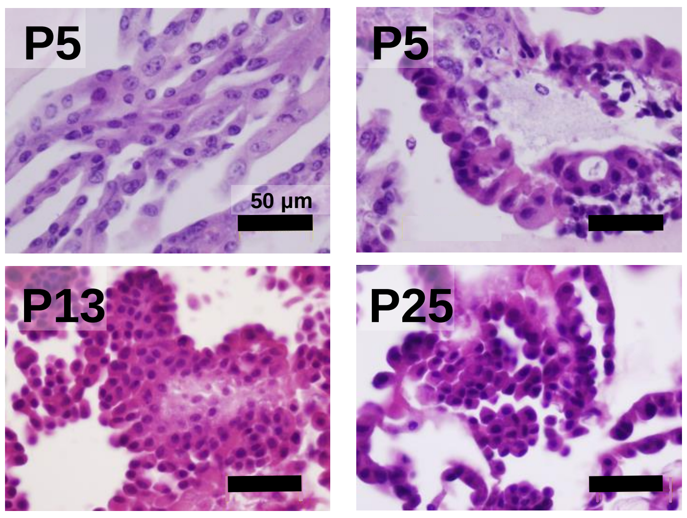


F

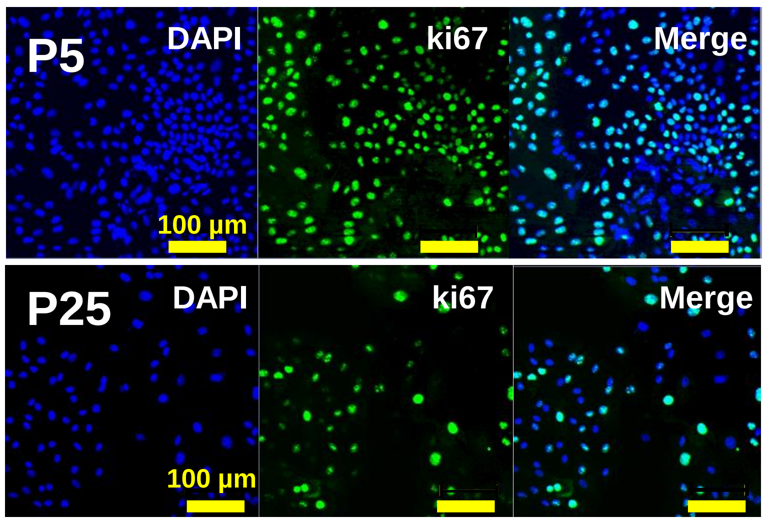


G

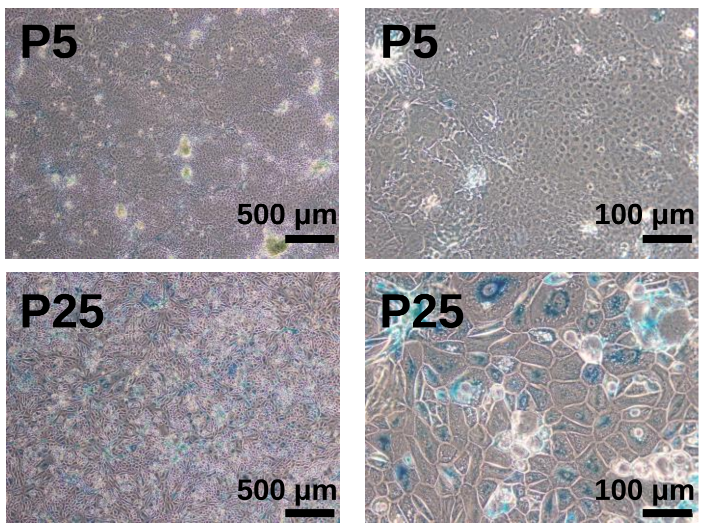


H

| Karyotype | Cell number |
|---|---|
| 46,XX | 14 |
| 45.X-X | 3 |
| 45,XX, dic(2;8) q37;p23) | 1 |
| 45,XX,der(8:17)(q10;q10),-16,-22 +2mar | 1 |
| 46,XX,del(17) (p11.1) | 1 |
| | 20 |

| Chromosome number | Cell number |
|---|---|
| 46 | 41 |
| 45 | 9 |
| | 50 |
